# Functional Connections Among Neurons within Single Columns of Macaque V1

**DOI:** 10.1101/2022.02.18.481095

**Authors:** Ethan B. Trepka, Shude Zhu, Ruobing Xia, Xiaomo Chen, Tirin Moore

**Affiliations:** Department of Neurobiology and Howard Hughes Medical Institute, Stanford University School of Medicine, Stanford, CA 94305, USA.; Department of Psychological and Brain Sciences, Dartmouth College, Hanover, NH 03755, USA.; Center for Neuroscience, Department of Neurobiology, Physiology, and Behavior, UC Davis, Davis, CA 95618, USA.

**Author notes:** These authors contributed equally to this work (Ethan B. Trepka, Shude Zhu).

## Abstract

Recent developments in high-density neurophysiological tools now make it possible to record from hundreds of single neurons within local, highly interconnected neural networks. Among the many advantages of such recordings is that they dramatically increase the quantity of identifiable, functional connections between neurons thereby providing an unprecedented view of local circuit interactions. Using high-density, Neuropixels recordings from single neocortical columns of primary visual cortex in nonhuman primates, we identified 1000s of functionally connected neuronal pairs using established crosscorrelation approaches. Our results reveal clear and systematic variations in the strength and synchrony of functional connections across the cortical column. Despite neurons residing within the same column, both measures of functional connectivity depended heavily on the vertical distance separating neuronal pairs, as well as on the similarity of stimulus tuning. In addition, we leveraged the statistical power afforded by the large numbers of connected pairs to categorize functional connections between neurons based on their crosscorrelation functions. These analyses identified distinct, putative classes of functional connections within the full population. These classes of functional connections were corroborated by their unique distributions across defined laminar compartments and were consistent with known properties of V1 cortical circuitry, such as the lead-lag relationship between simple and complex cells. Our results provide a clear proof-of-principle for the use of high-density neurophysiological recordings to assess circuit-level interactions within local neuronal networks.

## Introduction

Understanding the functional logic of local neuronal microcircuits is among the more fundamental objectives in the study of neural systems, yet it is also among the most challenging. This seems particularly true for mammalian neocortical circuits involved in perceptual and cognitive functions, and most notably in nonhuman primate model systems for which the available tools to interrogate those circuits are the most limited. The columnar organization of the mammalian neocortex (Lorente de No, 1938; Mountcastle, 1957) and its distinctly layered structure within different cortical domains are both widely appreciated (Reviewed in Defelipe, Markram, & Rockland, 2012; Horton & Adams, 2005; Mountcastle, 1997). In addition, several key principles of cortical circuitry, including constituent cell types (Harris & Mrsic-Flogel, 2013; Jiang et al., 2015; Katzel, Zemelman, Buetfering, Wolfel, & Miesenbock, 2011; Markram et al., 2004; Network, 2021; Packer & Yuste, 2011; Yoshimura & Callaway, 2005), input-output organization (Callaway, 1998; Douglas & Martin, 2004; Lefort, Tomm, Floyd Sarria, & Petersen, 2009; Munoz-Castaneda et al., 2021; Thomson & Bannister, 2003; Weiler, Wood, Yu, Solla, & Shepherd, 2008) and local microcircuit motifs (Avermann, Tomm, Mateo, Gerstner, & Petersen, 2012; Frandolig et al., 2019; Karnani et al., 2016; Obermayer et al., 2018; Pfeffer, Xue, He, Huang, & Scanziani, 2013; Pi et al., 2013) have emerged in recent years. Although it remains to be determined, such principles may turn out to generalize not only across neocortical areas, but also across species (Harris & Shepherd, 2015; Karten, 2015; Stacho et al., 2020) (but see Campagnola et al., 2021; Wildenberg et al., 2021). Yet, mapping complete cortical microcircuits within even a single cortical area remains a tremendous challenge (Adesnik & Naka, 2018).

Recent advances in recording technology have facilitated the development of large-scale, high-density micro-electrode arrays resulting in a substantial increment (>10x) in the number of neurons that can be studied simultaneously within a localized area of neural tissue. A prime example is the recent development of the Neuropixels probe (IMEC, Inc.), which consists of a high-channel count Si shank with continuous, dense, programmable recording sites (∼1000/cm). Numerous recent studies have demonstrated the advantages of such probes, such as their use in recording large neuronal populations within deep structures where optical approaches cannot be deployed (Jun et al., 2017; Steinmetz, Zatka-Haas, Carandini, & Harris, 2019). In addition, the high-density capacity of such probes dramatically increases the quantity of single neurons that can be obtained within a localized area of neural tissue (Siegle et al., 2021), thus making them well-suited for investigations of local neuronal circuitry. Given that studies of local neuronal circuitry within the primate brain are notoriously difficult to achieve, high-density electrophysiological approaches may be particularly valuable. However, only a few electrophysiological studies of the primate brain using such probes have been carried out thus far (Hesse & Tsao, 2020; Paulk et al., 2022; Sun et al., 2022; Trautmann et al., 2019; Zhu, Xia, Chen, & Moore, 2020). To date, many studies have leveraged the covariation in spiking activity between simultaneously recorded neurons to elucidate underlying neural mechanisms in the primate brain with some success, particularly within the visual system (Cohen & Kohn, 2011; Hansen, Chelaru, & Dragoi, 2012; Jia, Tanabe, & Kohn, 2013; Kohn & Smith, 2005; Koren, Andrei, Hu, Dragoi, & Obermayer, 2020; Smith & Kohn, 2008; Zandvakili & Kohn, 2015). In particular, temporally precise crosscorrelations in spiking activity have provided a unique means of assessing functional connectivity among neurons in both local and distributed networks (Aertsen & Gerstein, 1985; Aertsen, Gerstein, Habib, & Palm, 1989; Diba, Amarasingham, Mizuseki, & Buzsaki, 2014; Moore, Segundo, Perkel, & Levitan, 1970; Nelson, Salin, Munk, Arzi, & Bullier, 1992; Nowak, Munk, James, Girard, & Bullier, 1999; Perkel, Gerstein, & Moore, 1967; Siegle et al., 2021), and identification of functional connections has played an important part in understanding neural circuits in the mammalian visual system (Alonso & Martinez, 1998; Alonso, Usrey, & Reid, 1996, 2001; Baker & Bair, 2012; Briggs, Mangun, & Usrey, 2013; Hembrook-Short, Mock, Usrey, & Briggs, 2019; Michalski, Gerstein, Czarkowska, & Tarnecki, 1983; Reid & Alonso, 1995; Siegle et al., 2021; Toyama, Kimura, & Tanaka, 1981a; Ts’o, Gilbert, & Wiesel, 1986; Usrey, Reppas, & Reid, 1998, 1999). However, the extent of circuit-level details addressable with crosscorrelation is greatly limited by the low incidences of simultaneous recordings from connected neurons when using conventional extracellular recording techniques (e.g. Alonso et al., 2001; Hembrook-Short et al., 2019; Nelson et al., 1992; Ts’o et al., 1986). The use of high channel-count probes should substantially mitigate that limitation by virtue of the large increment in recording yield. Moreover, the high-density of recordings should further increase the incidence of isolating functionally connected neurons by virtue of the proximity of recorded cells.

We assessed the capacity of high-density Neuropixels probes to identify functionally connected neurons within cortical columns of primary visual cortex of macaque monkeys. Using established crosscorrelation approaches, we identified 1000s of functionally connected neuronal pairs during single recordings from neurons situated in different cortical layers. Our results demonstrate robust, systematic variations in the strength and synchrony of functional connections across the cortical column. In addition, by leveraging the large numbers of connected pairs, distinct classes of functional connections could be identified within the full population.

## Results

### Identifying functional connections within single columns of visual cortex

The activity of V1 neurons was recorded in two anesthetized macaque monkeys (M1, M2) using high-density, multi-contact Neuropixel probes (version 3A; IMEC Inc, Belgium) (Fig. 1a) (**Methods**). Each probe consisted of 986 contacts (12 mm x 12 mm, 20 µm spacing) distributed across 10 mm, of which 384 contacts could be simultaneously selected for recording. Probes were inserted into the lateral operculum of V1 with the aid of a surgical microscope at angles nearly perpendicular to the cortical surface. The dense spacing between electrode contacts provided multiple measurements of the waveforms from individual neurons (mean = 4.52 measurements) (Fig 1a, b) and facilitated the isolation of large numbers of single neurons. In each of 5 experimental sessions (3 in M1, 2 in M2), we measured the visual responses of 115-221 simultaneously recorded neurons to drifting gratings presented at varying orientations (total = 802 neurons). As expected, neurons were highly orientation selective, and exhibited both simple and complex cell properties (De Valois, Albrecht, & Thorell, 1982; Hubel & Wiesel, 1962, 1968) (Fig. 1b). As in previous studies (Briggs et al., 2013; Hembrook-Short et al., 2019; Jia et al., 2013; Kohn & Smith, 2005; Siegle et al., 2021; Smith & Kohn, 2008; Zandvakili & Kohn, 2015), we used the visually driven spike trains to measure crosscorrelations between simultaneously recorded neuronal pairs.

**Figure 1.**
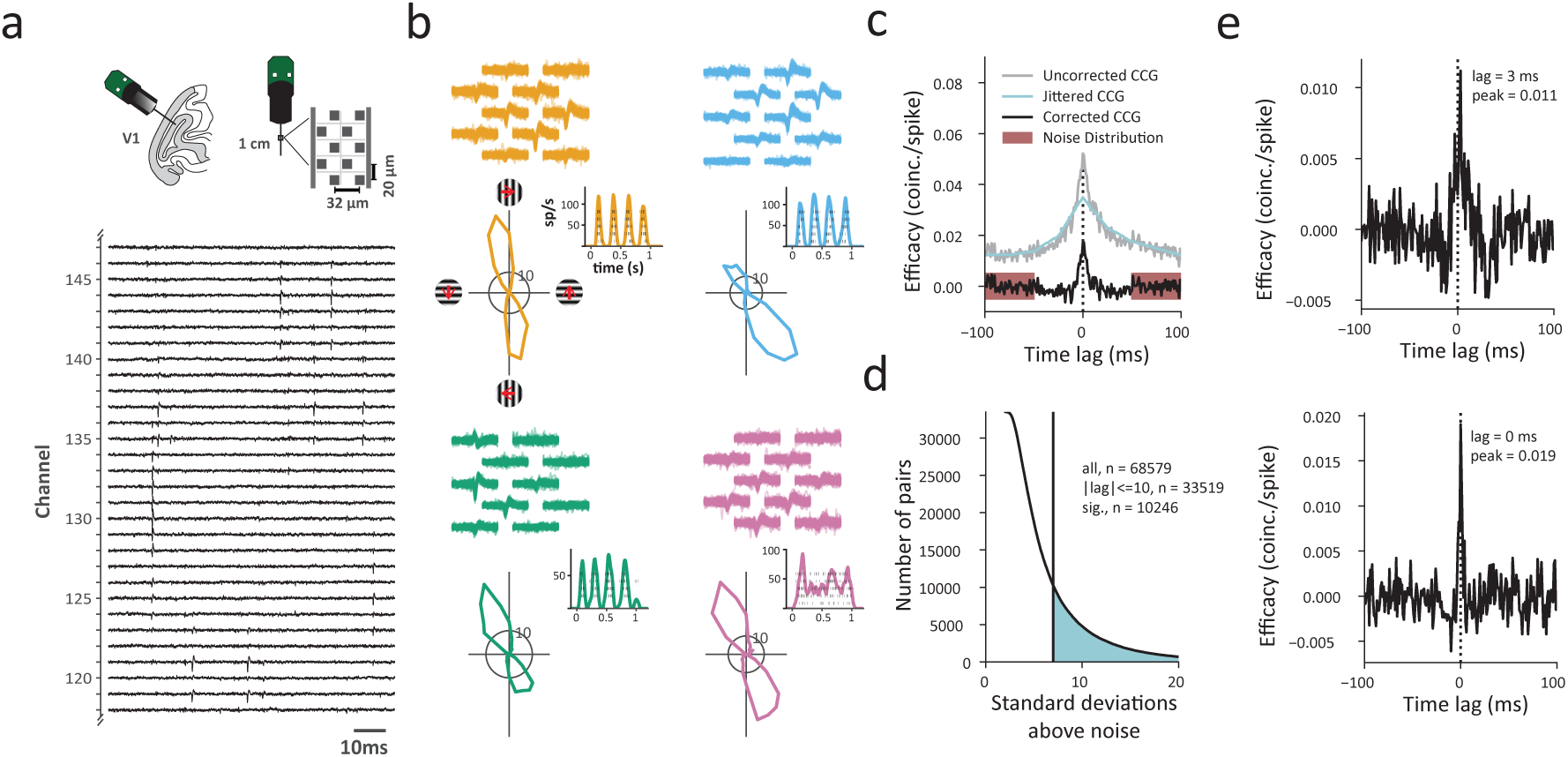
Identifying functional connections within single columns of visual cortex. **a,** Upper left cartoon depicts the angle of Neuropixel probe penetrations made into the lateral surface and underlying calcarine sulcus of V1. Upper right, Neuropixels probe base and shank, and layout of electrode contacts for a section of the recording shank. Lower, raw voltage traces recorded from an exemplar section of channels and time period. **b**, Example single-neuron recordings with Neuropixels probes, three simple cells (orange, blue, green) and one complex cell (purple). Top, spike waveforms recorded across multiple adjacent electrode contacts are shown for each neuron. Bottom, each neuron’s response to its preferred orientation (rasters and instantaneous spike rates) and their corresponding tuning curves. Red arrows (uppler left) denote the drift direction of oriented gratings. **c,** Example CCG between an example pair of V1 neurons. Corrected CCGs were generated from the difference between a jittered and an uncorrected CCG. Significance of each CCG was determined from comparisons between the peak and the noise distribution. **d,** Distribution of ratios of CCG peaks to the noise (SD) for all recorded pairs. Shaded area denotes CCGs with peaks >7 SDs above the mean of the noise distribution. **e,** Two example CCGs differing in both peak lag and peak efficacy.

To estimate the functional connectivity between pairs of neurons recorded simultaneously within columns of V1, we computed cross-correlograms (CCGs) using the 802 visually responsive neurons recorded across sessions. CCGs were computed from the spike trains of 68,579 pairs of simultaneously recorded neurons (6,555 – 24,310 pairs/session) (**Methods**). Each CCG was normalized by the firing rate (FR) and jitter-corrected to mitigate the influences of FR (Bair, Zohary, & Newsome, 2001; Mastronarde, 1983) and correlated slow fluctuations (Harrison & Geman, 2009; Smith & Kohn, 2008), respectively, yielding a corrected CCG (Fig. 1c). In addition, as in previous studies, we considered a CCG significant only if its peak occurred within 10 ms of zero time lag, and if that peak was > 7 standard deviations above the mean of the noise distribution (Siegle et al., 2021). Using this criterion, a total of 10,246 significant CCGs were obtained from all recording sessions (Fig. 1d), with each session yielding 755-3,022 significant CCGs. The peak lag of each CCG, defined as the differences between zero and the time when the peak occurred, estimates the synchrony and/or direction of functional connectivity between neuronal pairs; whereas the peak efficacy measures the strength the connectivity (Fig 1e).

### Variation in the synchrony and strength of functional connections within cortical columns

A number of previous studies using low-channel count probes or chronically implanted electrode arrays have shown that correlated activity in primate V1 declines with the horizontal distance separating pairs of neurons (Kruger & Aiple, 1988; Maldonado, Friedman-Hill, & Gray, 2000; Smith & Kohn, 2008) (see also Chu, Chien, & Hung, 2014). Evidence from these studies suggest that correlations are greatest for pairs of neurons located within the same column, and diminish with greater columnar distance. Other evidence shows variation in the spike timing correlations between neuronal pairs located within different laminar compartments (Smith, Jia, Zandvakili, & Kohn, 2013). However, considerably less is known about how the nature of correlations varies across the depth of individual columns where the degree of shared input and connectivity is at its highest. We therefore leveraged the large numbers of significantly correlated pairs obtained from high-density recordings to examine how the synchrony and strength of correlations depended on the vertical distance separating neurons within V1 columns. Figure 2a shows data from an example recording session in which 221 visually responsive neurons were recorded and 2,453 significantly correlated pairs were obtained. All neurons are shown along the ∼2 mm depth of cortex. Shown also are two example correlated pairs whose CCGs are shown in Figure 1e. Of the two pairs, the vertical distance separating neurons in one pair was 138 µm greater than that of the other. In spite of this small difference, CCG of the closer pair was both more synchronous and stronger than the more distant pair. This pattern of results was observed across all significantly correlated pairs and in all sessions (Fig. 2b-c) (Figs S1-S2). The synchrony of correlated spiking diminished several fold across neuronal pair distance. This change could be fit with a linear function (r = 0.42; P<10^-5^) in which the (absolute) peak lag increased at a rate of 1.3 ms / 500 µm of vertical distance. Peak efficacy of the significant CCGs also depended heavily on pair distance. This effect could be fit with an exponential decay function (r = -0.34; P<10^-5^) in which the peak efficacy decreased by half within 154 µm. Thus, both measures of functional connectivity depended heavily on the vertical distance separating neuronal pairs. In addition, we confirmed that the effects of vertical distance on both the synchrony and strength were independent of whether neuronal pairs were located within the same or different cortical layers (Fig. S3).

**Figure 2.**
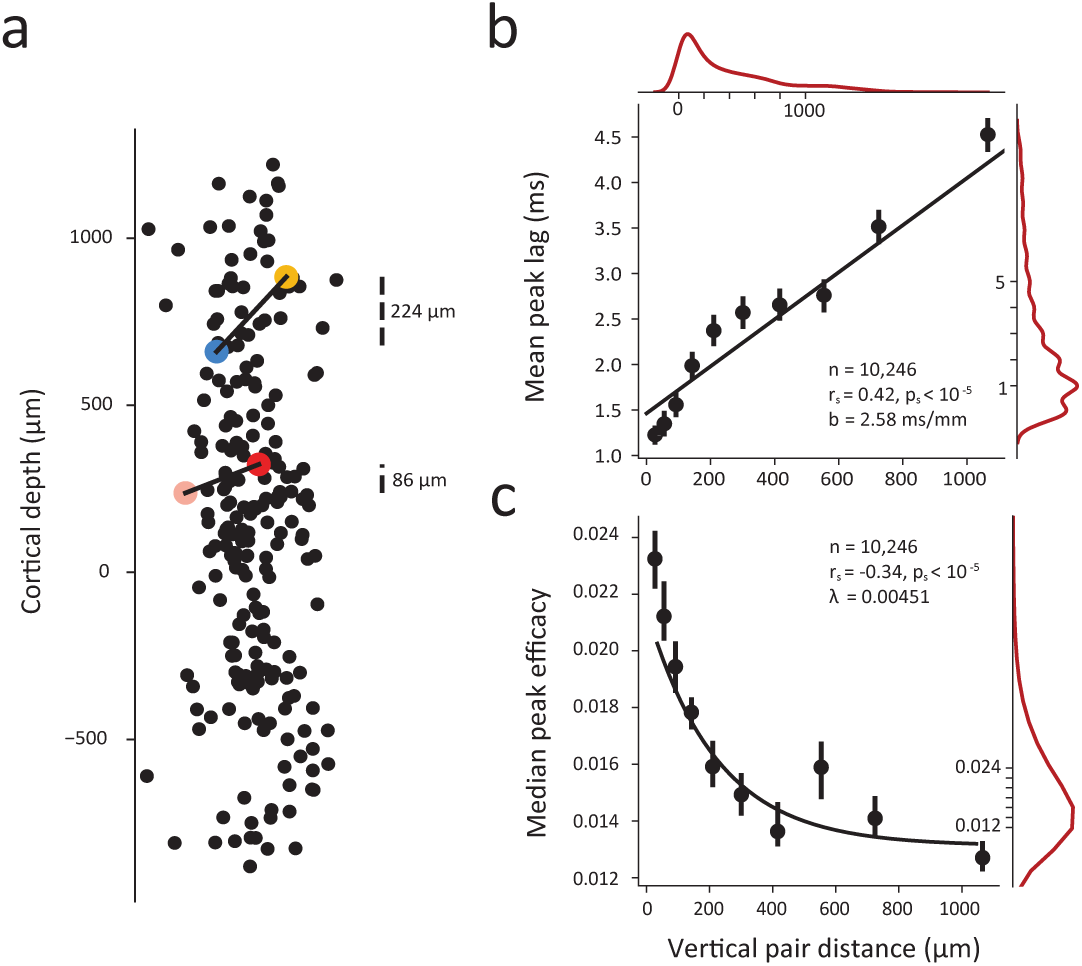
Dependence of synchrony and strength of functional connectivity on vertical distance within single cortical columns. **a,** Example session (M1, session 3) with 221 visually responsive neurons recorded simultaneously and their locations across cortical depth. (Horizontal axis is magnified for visualization). Cortical depth 0 denotes the boundary between Layer 4C and Layer 5. Laminar boundaries were determined using histological data and current-source-density (CSD) profile for each session (Methods). Two example correlated pairs from Figure 1e with varied CCGs are shown in color (blue-yellow pair and pink-red pair corresponding to Figure 1e top and bottom, respectively). **b**, Linear dependence of synchrony on vertical pair distance. **c, S**trength of CCGs decay with greater pair distance. In b and c, all significantly correlated pairs from all sessions are combined and each dot denotes the mean peak lag or median peak efficacy of significantly correlated CCGs within a (10% quantile) vertical distance bin. Error bars denote 95% confidence intervals. Black lines denote the linear and exponential fits in b and c, respectively; slope (b) and decay constant (λ) are shown. Red lines denote marginal distributions.

### Dependence of synchrony and strength of functional connections on tuning similarity

In addition to the dependence of correlated activity on the distance between neuronal pairs, many studies have shown that greater functional and synaptic connectivity typically occurs between neurons with similar stimulus preferences (Chu et al., 2014; Constantinidis, Franowicz, & Goldman-Rakic, 2001; Cossell et al., 2015; DeAngelis, Ghose, Ohzawa, & Freeman, 1999; Denman & Contreras, 2014; Funahashi & Inoue, 2000; W. C. Lee et al., 2016; Ts’o et al., 1986) (but see Das & Gilbert, 1999; Maldonado et al., 2000). Within primate V1, stimulus selectivity is notably similar for neurons within the same column, particularly for orientation selectivity (Blasdel & Salama, 1986; Hubel & Wiesel, 1968, 1974; Ts’o, Frostig, Lieke, & Grinvald, 1990), and this was evident in our recording sessions, where the peak visual responses were largely aligned at the same stimulus orientation across cortical depth (Fig. 3a). We considered that within orientation columns, functional connectivity could be homogenous for populations of similarly tuned neurons. Alternatively, it could be that even small variations in tuning similarity could result in robust differences in the synchrony and strength of correlated activity. To address this, we examined the dependence of synchrony and strength on the similarity of visual properties of neurons within the same cortical column. As in previous studies (Shadlen & Newsome, 1998; Zohary, Shadlen, & Newsome, 1994), we quantified tuning similarity by computing signal correlations (r_ori_) for each neuronal pair (**Methods**). Across the total number of neuronal pairs (N = 68,579), the mean r_ori_ was 0.25. For the significantly correlated neuronal pairs, the mean r_ori_ was 0.33. Signal correlations for the two previous example neuronal pairs are shown in Fig 3b. The responses of both pairs are positively correlated, yet that correlation is much higher in the second, more proximal, pair (Fig. 2a) and the one with more synchronous and stronger CCG (Fig 1e). Overall, we found that both the peak lag and peak efficacy of CCGs for significantly correlated neuronal pairs varied monotonically with tuning similarity across the range of signal correlations (Fig. 3c, d). Neuronal pairs with the highest signal correlations exhibited half the peak lags and twice the peak efficacies of uncorrelated pairs. This pattern was observed in each of the individual recording sessions (Figs S4-S5).

**Figure 3.**
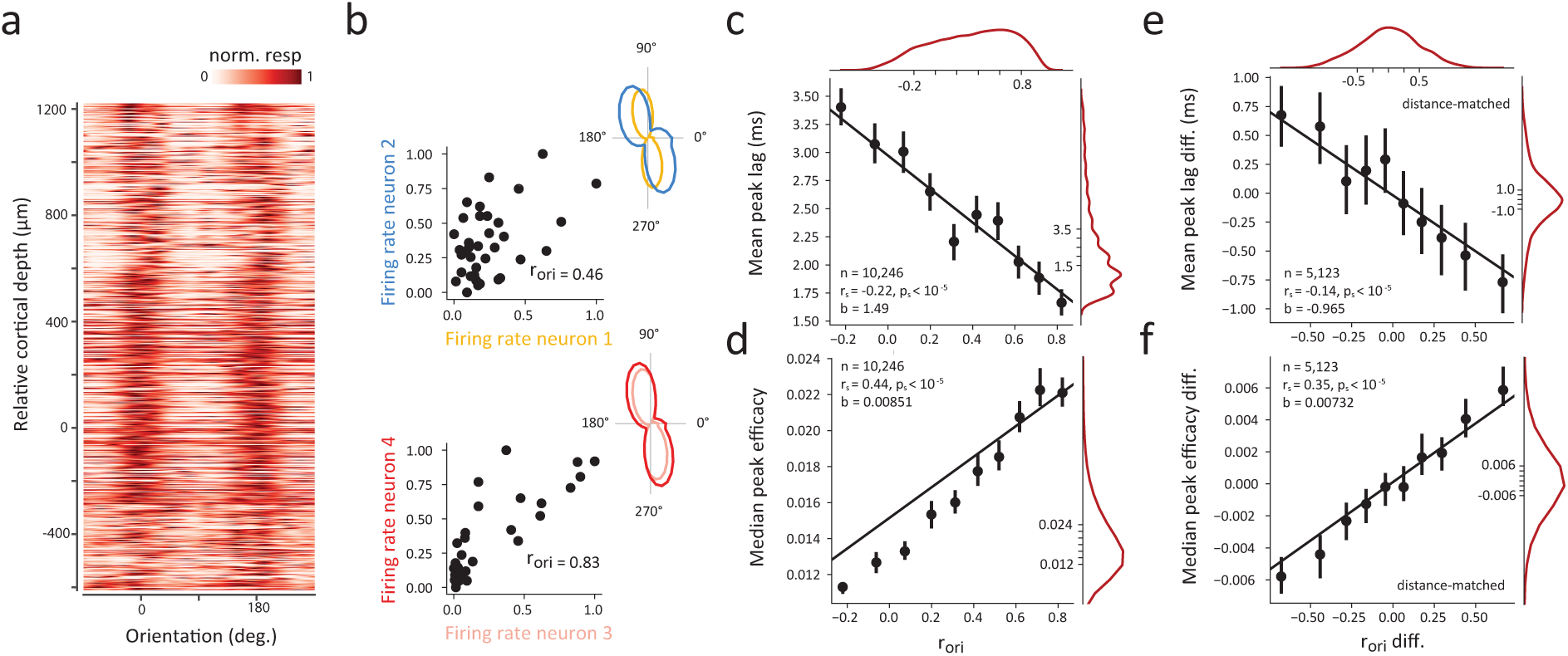
Dependence of synchrony and strength of functional connectivity on tuning similarity within single cortical columns. **a,** Heat map of visual responses across drift directions of oriented gratings and across cortical depth. The response tuning of each of 802 neurons was aligned to the overall preferred orientation shared by neurons recorded from the same session, and all sessions were combined. **b**, Signal correlation between exemplar neurons. Left, Scatter plot of normalized responses to different stimulus orientations (n=36) for the two example pairs shown in Figure 1e and Figure 2a. Signal correlations (r_ori_) are also shown. Right, each neuron’s orientation tuning curve. **c**, Linear dependence of synchrony on the corresponding signal correlation. **d,** Linear dependence of CCG strength on the corresponding signal correlation. **e**, Difference in peak lag of distance-matched CCGs was positively correlated with difference in signal correlation. **f**, Difference in peak efficacy of distance-matched CCGs was negatively correlated with difference in signal correlation. In c-f, all significantly correlated pairs from all sessions are combined and each dot denotes mean peak lag or median peak efficacy of significantly correlated CCGs within a (10% quantile) signal correlation bin. Error bars denote 95% confidence intervals. Black lines denote linear fits; slopes (b) are shown. Red lines denote marginal distributions.

We considered that the apparent relationship between the synchrony and strength of functional connections and signal correlation might result indirectly from a collinear effect of vertical distance on CCGs (Fig. 2). To address this, we examined differences in the peak lags and peak efficacies of CCGs between combinations of two neuronal pairs separated by comparable cortical distances. Specifically, we sorted all significantly correlated CCGs by their vertical distances, and then examined whether differences in signal correlations (r_ori_) among adjacently sorted (distance-matched) pairs, were still associated with differences in CCG peak lags and peak efficacies (**Methods**). Indeed, we found that the differences in peak lags of distance-matched CCGs were positively correlated with signal correlation (Fig. 3e) and the differences in peak efficacies of distance-matched CCGs were negatively correlated with signal correlation (Fig. 3f). These results indicate that signal correlations within the column predicted both the synchrony and strength of functional connections independent of vertical pair distance. Nonetheless, the distance-matched correlations (Fig. 3e-f) were smaller than their corresponding unmatched correlations (Fig. 3c-d), suggesting that the vertical distance between neurons and their orientation signal correlations exhibit distinct, but overlapping, effects on the strength and timing of functional connections within a single cortical column.

To quantify the distinct contributions of vertical pair distance and orientation signal correlation to the synchrony and strength of CCGs, we fit GLMs to predict CCG peak lag and peak efficacy using pair distance and signal correlation as predictors (**Methods**). Predictors were standardized so their relative effects could be compared, and peak outliers (1.5*IQR criterion) were removed. The resulting regression equations were:

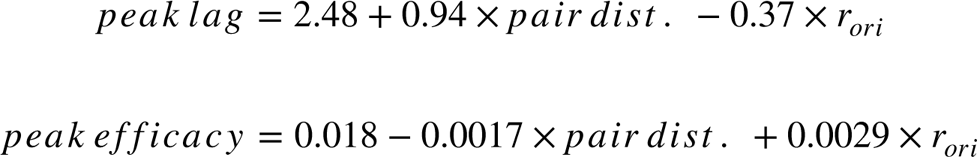

Regressions explained 19% of variance in peak lag (*R*^2^ = 0.191) and 20% of variance in peak efficacy (*R*^2^ = 0.195). In the regression predicting CCG peak lag, the coefficient of pair distance was nearly three times the coefficient of signal correlation. In contrast, for CCG peak efficacy, the coefficient of signal correlation was nearly twice that of pair distance. Furthermore, whereas signal correlation was less predictive of CCG peak lag, it was more predictive of CCG peak efficacy than pair distance.

### Classification of functional connections

CCG peak lags and peak efficacies are often the parameters of interest in crosscorrelations (Briggs et al., 2013; Hembrook-Short et al., 2019; Smith & Kohn, 2008), yet they are simplifications of the more complex, underlying crosscorrelation functions. The shape of these correlation functions may offer additional insights into the distinct types and properties of functional connections present among neurons within a network. Several theoretical studies have suggested a correspondence between CCG shape and underlying pairwise connectivity (Aertsen & Gerstein, 1985; Melssen & Epping, 1987) that can be further influenced by overall network structure and background noise (Ostojic, Brunel, & Hakim, 2009). For example, synchronous CCGs tend to correspond to pairs of neurons that receive input from a common source, while asynchronous CCGs tend to correspond to pairs that have direct synaptic connections (Ostojic et al., 2009). Moreover, synchronous CCGs with narrow peaks and synchronous CCGs with broad peaks may correspond to pairs of neurons that receive input from common sources with shorter and longer autocorrelation timescales, respectively (Ostojic et al., 2009). Experimental studies have corroborated these findings, identifying similar CCG shapes in different cortical regions and species (Alonso & Martinez, 1998; Constantinidis et al., 2001; Hembrook-Short et al., 2019; Siegle et al., 2021). However, the distribution of these CCG shapes within a single cortical column remains unknown. Furthermore, whether that distribution within V1 corroborates other evidence about the functional and/or anatomical relationships among V1 laminae and cell types remains unclear.

To address these questions, we clustered the entire population of CCGs, taking advantage of the large number of connected pairs to identify robust CCG templates. To do this, we first normalized significant CCGs, and utilized t-distributed Stochastic Neighbor Embedding (t-SNE) to map CCGs to a lower-dimensional space, and then clustered CCGs in the resulting space using k-means (**Methods**). To select a statistically reasonable number of clusters, we examined how the total variance and the silhouette score explained by clustering changed as a function of the number of clusters (Fig. 4a). From this, we selected four as the optimal number of clusters given that silhouette score peaked ∼3-4 clusters, and 4 clusters explained more variance than 3.

CCG shape was relatively heterogenous within each of the four clusters (Fig. 4b). Nonetheless, by averaging over all CCGs in each cluster, we could construct CCG templates that summarized key characteristics of the clusters (Fig. 4c). Within the full population, we identified two synchronous classes of functional connections, a ‘sharply synchronous’ class (*S_sync_*) with an narrow peak at *τ* = 0 and a ‘broadly synchronous’ class (*B_sync_*) with a wide peak at *τ* = 0. In addition, two asynchronous classes were identified, a ‘forward’ class (*F_async_*) (leading) and a ‘reverse’ class (*R_async_*) (lagging) with more probability density before and after *τ* = 0 (median *τ 3ms*), respectively (Fig. 4c). Aside from clear differences in peak lags between subsets of the putative classes (e.g. synchronous vs. asynchronous), CCGs of different classes also differed in their peak efficacies; synchronous classes exhibited higher average peak efficacies than asynchronous classes (median peak efficacy: *S_sync_* 0.021, *B_sync_* 0.020, *F_async_* 0.015, *R_async_* 0.014). Importantly, our objective was not to find the exact number of distinct classes of functional connections in V1 or to perfectly categorize every functional connection into a homogenous cluster. Instead, we sought to identify at least one set of clusters that is consistent with that expected in local microcircuits

### Corroboration of putative CCG classes with V1 microcircuitry

We next examined the extent to which the putative CCG classes were also distinguishable from one another along anatomical and functional lines given other known properties of V1 microcircuits. First, we considered that the identified classes might differ in their vertical pair distances and signal correlations. Indeed, we found that vertical pair distances were larger and orientation signal correlations were smaller in asynchronous (*F_async_* and *R_async_*) than in synchronous (*S_sync_* and *B_sync_*) classes (Fig 5a-b) (significant pairwise comparisons: p<10^-5^). Although, this result is expected given that both the peak lag and peak efficacy components of CCGs were clearly predicted by distance and signal correlation (Figs. 2-3), additional differences emerged between the identified classes. For example, we found that in spite of similar CCG peak lags, *B_sync_* pairs were separated by greater vertical distances than *S_sync_* ones (Fig. 5a) (p<10^-5^). Furthermore, in spite of being separated by a greater cortical distance, *B_sync_* pairs exhibited higher signal correlations than *S_sync_* pairs (Fig 5b) (p<10^-5^). These findings thus provide some validation of the apparent subtypes of CCGs.

**Figure 4.**
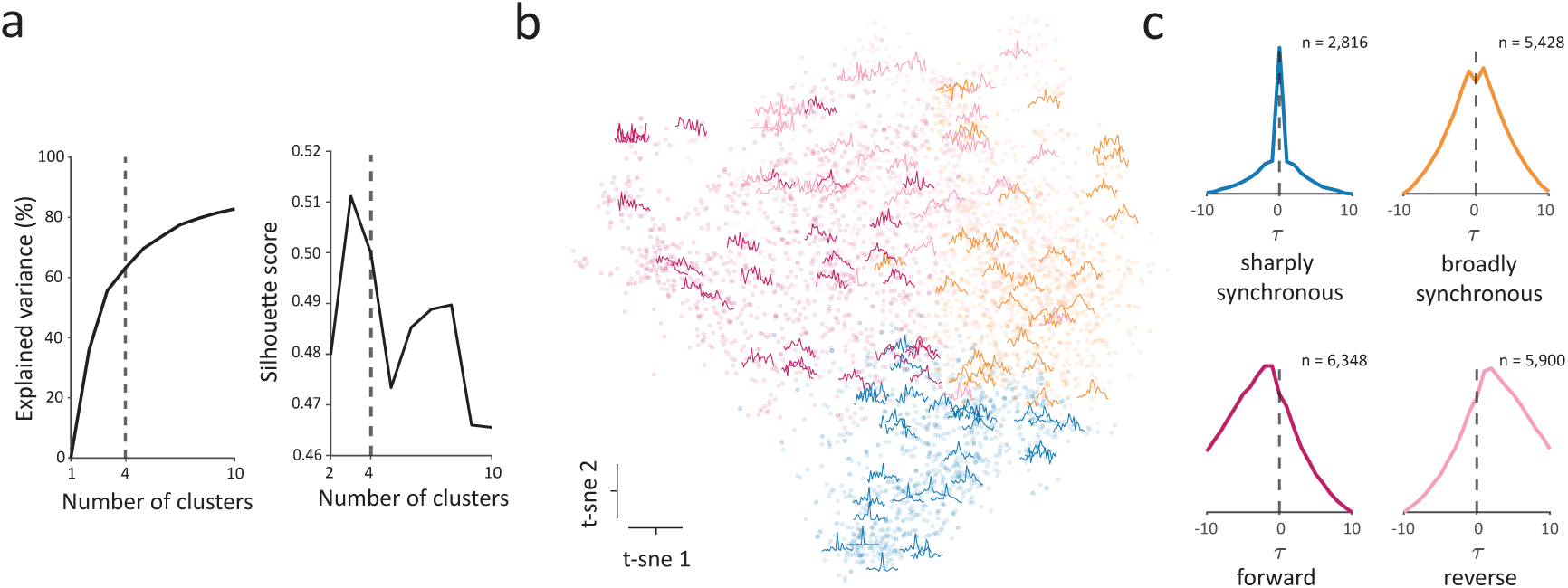
Identification of distinct classes of functional connections within the full population. **a**, Explained variance (left) and silhouette score (right) as a function of number of clusters. The dashed vertical line indicates the selected number of clusters (n=4). **b,** Scatter plot of dimensionality-reduced CCGs in the first two dimensions of t-SNE space. Randomly selected example CCGs are overlayed on the scatterplot in their corresponding location in t-SNE space. **c**, CCG templates generated by averaging over all the CCGs in each cluster. The templates include a ‘sharply synchronous’ class (*S_sync_*) with a narrow peak at *τ* = 0, a ‘broadly synchronous’ class (*B_sync_*) with a wide peak at *τ* = 0, a ‘forward’ class (*F_async_*) (leading) with more probability density before *τ* = 0 and a ‘reverse’ class (*R_async_*) (lagging) with more probability density after *τ* = 0.

**Figure 5.**
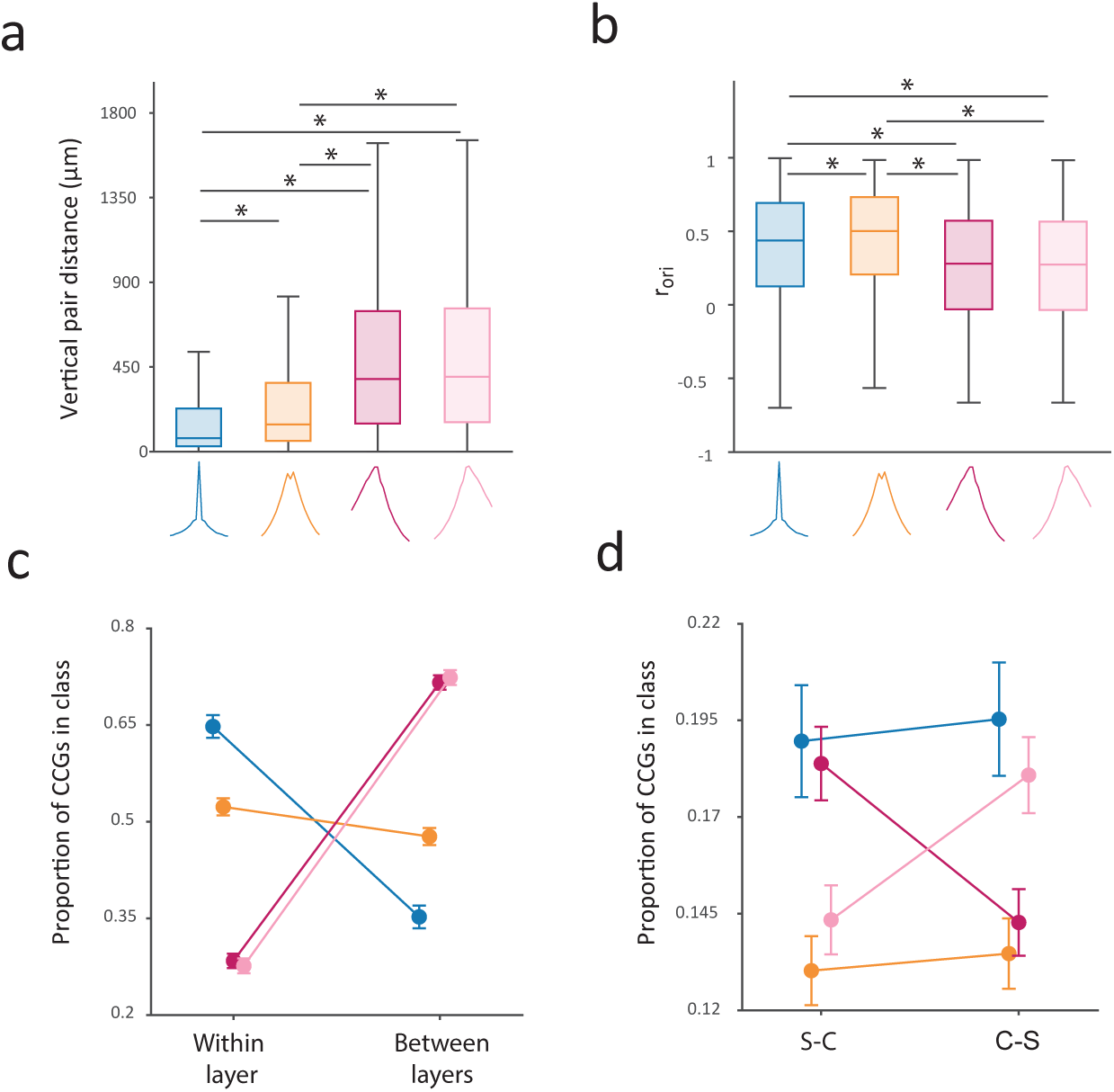
Corroboration of putative classes of functional connections with V1 microcircuitry. **a,b,** Boxplots of vertical pair distance (a) and orientation signal correlation (b) across the 4 identified CCG classes. Boxplots illustrate the medians, first and third quartiles, and non-outlier (1.5*IQR method) minima and maxima. Asterisks denote significant differences in medians between pairs of classes (Wilcoxon rank-sum test; p<10^-5^). **c**, Proportion of CCGs in each class composed of two neurons from the same (‘within layer’) or different layers (‘between layer’). **d**, Proportion of CCGs in each class composed of a simple cell and a complex cell computed with either the simple cell (S-C) or the complex cell (C-S) as the first neuron in the crosscorrelation function. In c and d, error bars indicate margin of error with a 95% confidence interval.

Next, we examined whether the identified classes of functional connections differed in their laminar distributions. Indeed, we found that different classes were differentially distributed across V1 layers such that one or more of the identified classes were often overrepresented among functional connections within particular layers (Fig. S6) (p<10^-5^). To simplify this result, we compared the proportion of CCGs in each class composed of two neurons in the same layer or different layers. We found that most of the asynchronous pairs were composed of neurons from different layers, while most of the synchronous pairs, particularly the *S_sync_* ones, were composed of neurons from the same layer (Fig 5c) (within vs between proportion: *S_sync_* [0.65 vs 0.35], *B_sync_* [0.52 vs 0.48], *F* [0.28 vs 0.72], *R* [0.28 vs 0.72], one proportion z-test: p<10^-3^). This observation dovetails the relationship between pair distance and CCG class described above. Nonetheless, we found that cortical layer had an independent effect of distance on CCG class assignment among nearby pairs of neurons. The laminar composition of functional connections (within vs between) was a significant predictor if CCGs were members of the *B_sync_* class, but not the *S_sync_*, *F_async_*, or *R_async_* classes when controlling for the effects of vertical distance (Table S2; logistic regression, p<10^-5^). More specifically, CCGs composed of two neurons within the same layer had a higher probability of falling in the *B_sync_* class than CCGs with comparable vertical distances composed of two neurons in different cortical layers.

In addition to its laminar organization, V1 neurons exhibit clear differences in their receptive field properties. In particular, V1 neurons classically fall into two broad functional types: simple (S) and complex (C) cells (De Valois et al., 1982; Hubel & Wiesel, 1962, 1968; Movshon, Thompson, & Tolhurst, 1978; Skottun et al., 1991) (see also Chance, Nelson, & Abbott, 1999; Mechler & Ringach, 2002; Priebe, Mechler, Carandini, & Ferster, 2004). Complex cells appear to receive converging input from groups of simple cells, and this fact suggests that simple cells should lead rather than lag complex cells in their CCGs. To test this in our data, we compared the proportion of significant CCGs in each class composed of one simple and one complex cell. Among the significant CCGs, a majority were comprised of pairs of complex cells (S-S=0.052, C-C=0.630; S-C or C-S=0.318; one-proportion z-test: p<10^-5^). Among the mixed (S-C or C-S) pairs, CCGs were computed with either functional type as the first or second neuron in the crosscorrelation function, i.e., S-C or C-S (**Methods**). Thus, we could compare the proportions of those two subsets among the different CCG classes (Fig. 5d). We found that the proportion of ‘forward’ (*F_async_*) CCGs was larger when simple cells were chosen as the first neuron than when complex cells were chosen as the first neuron. Likewise, the proportion of ‘reverse’ (*R_async_*) CCGs was larger when instead complex cells were chosen as the first neuron than when simple cells were chosen as the first neuron (chi-squared test; p<10^-5^). Notably, although the dominant lead-lag relationship between simple and complex cells is consistent with established models of V1 (Alonso & Martinez, 1998; Martinez & Alonso, 2001; Yu & Ferster, 2013), there were also many CCG pairs in which complex cells led simple cells or where the pair fired synchronously. This heterogeneity in functional interactions between simple and complex cells is consistent with studies suggesting that simple and complex cells might arise from variations in a continuous process as opposed to being two clearly distinct populations (Chance et al., 1999; Kim, Jang, & Paik, 2021; Mechler & Ringach, 2002; Priebe et al., 2004).

### Corroboration of different classes with V1 input and local circuitry

Previous studies have characterized the anatomical organization of dorsal lateral geniculate nucleus (dLGN) input to V1 in extensive detail (Blasdel & Lund, 1983; Hendrickson, Wilson, & Ogren, 1978; Hubel & Wiesel, 1972). In the macaque brain, dLGN magnocellular and parvocellular axons primarily project to V1 layers 4cα and 4*cβ*, respectively, along with inputs that terminate in layer 6 (reviewed in Briggs & Usrey, 2011; J. S. Lund, 1988; Merigan & Maunsell, 1993; Nassi & Callaway, 2009) (Fig. 6a). However, the extent to which functional connections within layers of V1 reflect these anatomical projections remains unclear. Thus, we examined the distribution of CCG classes across pairs of V1 input layers 4c*α*, 4*cβ*, and 6. We found that for the 4cα-4c*α*, 4*cβ*-4*cβ*, and 6-6 pairings, *S_sync_* CCGs were observed much more frequently than other CCG classes (Fig. 6b) (chi-squared test; p<10^-5^). This overrepresentation of *S_sync_* CCGs may reflect the fact that neurons in 4cα, 4*cβ*, and 6 receive common and converging input from the dLGN. Furthermore, it is noteworthy that the *S_sync_* class was overrepresented in V1 input layers but the *B_sync_* class was not.

**Figure 6.**
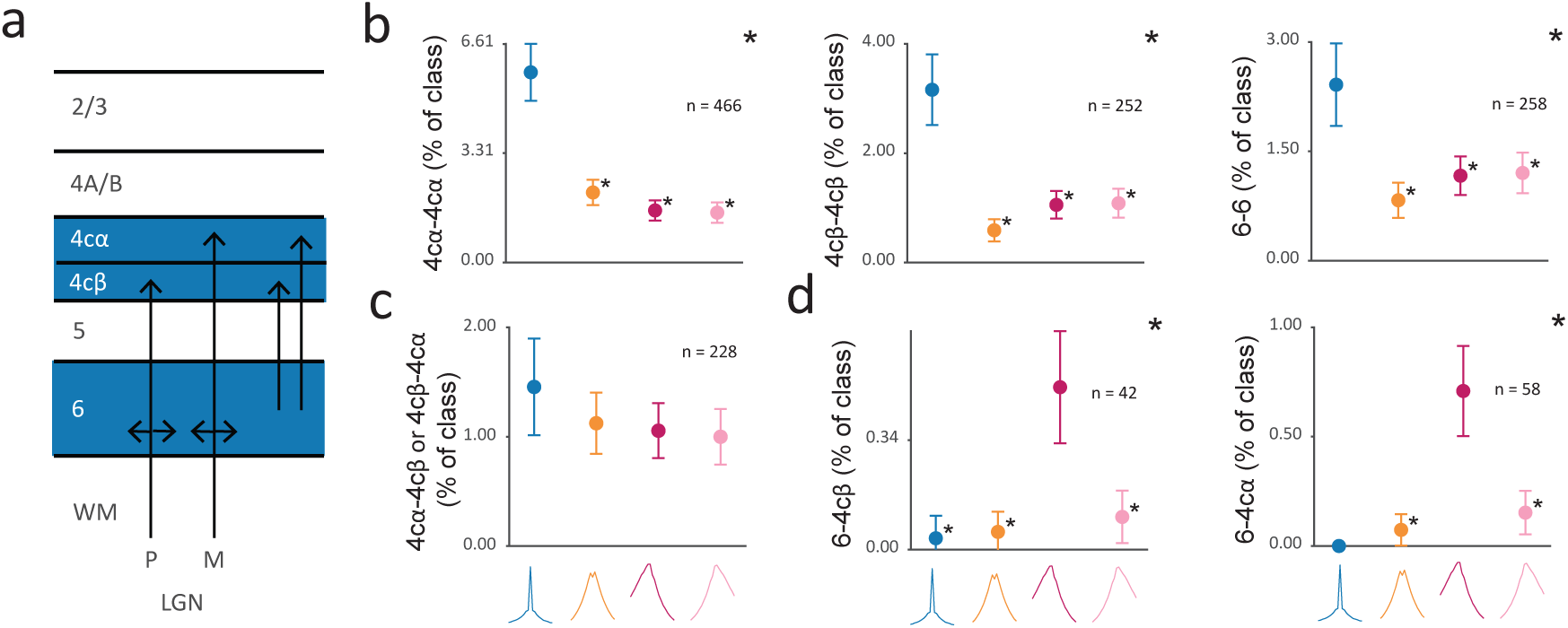
Distribution of different putative classes of functional connections within V1 input layers. **a**, Diagram of dLGN input to V1 layers 4cα, 4c*β*, and 6; dLGN axons terminate in layers 4cα and 4*cβ* and layer 6, and layer 6 projects to layers 4cα and 4c*β*. **b,** Percentage of CCGs in each class composed of two neurons in layer pairings of 4cα-4cα, 4c*β*-4c*β*, or 6-6, out of all the pairwise layer pairing combinations. **c**, Percentage of CCGs in each class composed of one neuron in layer 4cα and one in 4c*β* (4c -4c*β* or 4c*β*-4c). **d**, Percentage of CCGs in each class composed of one neuron in layer 6 and the other in 4c*β* (left) and 4cα (right) computed with the layer 6 neuron as the first neuron in the crosscorrelation function (6-4c*β*, 6-4cα). For b-d, error bars denote 95% confidence intervals. Large asterisks in the upper right denote significant chi-squared test of all classes versus the selected layer pairing (Bonferonni correction for 60 comparisons, 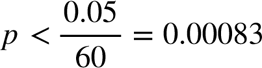). Small asterisks near each dot denote significant chi-squared test of layer pairing versus the class with asterisk and the class with maximum proportion in the panel ( *p* < 0.00083).

In contrast to the overrepresentation of *S_sync_* CCGs within the input layers, this class of CCGs was not overrepresented in functional connections between input layers. Of the four CCG classes, the proportions of each found among pairs composed of one neuron in layer 4cα and one neuron in layer 4*cβ* were statistically indistinguishable (Fig. 6c) (chi-squared test; *p* = 0.27). The lack of an overrepresentation of *S_sync_* CCGs among 4c*α* − 4*cβ* pairs could reflect the lack of synchrony between magnocellular (fast) and parvocellular (slow) inputs to V1. This result is noteworthy given that the average distance between neuronal pairs across 4cα and 4*cβ* was comparable to the distances between neuronal pairs within V1 input layer 6 (mean distance: 4cα - 4*cβ* = 106 µm; 6-6 = 80 µm). In examining functional connections between layers 4Cα and 6, we considered that a temporal offset between layer 6 and 4Cα neurons might exist given extensive projections from layer 6 pyramidal neurons to layer 4C (Wiser & Callaway, 1996) To test this, we examined the 6-4c and 6-4*cβ* subsets of pairs in which the layer 6 neuron was the first neuron in the crosscorrelation function. Indeed, in addition to observing that *S_sync_* CCGs were poorly represented, we found that the *F_async_* class was significantly overrepresented in these pairs (Fig. 6d) (p < 10^-5^).

Lastly, we examined the distribution of CCG classes across pairs of neurons involving layer 2/3 neurons. A wealth of evidence indicates that layer 2/3 neurons provide a major source of output to other neocortical areas (reviewed in Callaway, 1998; Douglas & Martin, 2004; Felleman & Van Essen, 1991; Harris & Shepherd, 2015; Thomson & Lamy, 2007)). In macaque V1, layer 2/3 neurons send projections to higher visual areas such as V2 (Livingstone & Hubel, 1984; Rockland, 1992; Sincich & Horton, 2005) and V4 (Yukie & Iwai, 1985), and receive inputs from all the deeper cortical layers, including layer 4C, 4C*β*, 4A, 4B, 5 and 6 (Blasdel, Lund, & Fitzpatrick, 1985; Callaway, 1998; Callaway & Wiser, 1996; Fitzpatrick, Lund, & Blasdel, 1985; Kisvarday, Cowey, Smith, & Somogyi, 1989; Lachica, Beck, & Casagrande, 1992; Jennifer S Lund & Boothe, 1975; Sawatari & Callaway, 2000; Vanni, Hokkanen, Werner, & Angelucci, 2020; Wiser & Callaway, 1996; Yarch, Federer, & Angelucci, 2017; Yoshioka, Levitt, & Lund, 1994) (Fig. 7a). Consequently, one might predict that a predominant proportion of projections to 2/3 neurons from other layers might be forward ones (Callaway, 1998; Mejias, Murray, Kennedy, & Wang, 2016; Schmidt et al., 2018). Consistent with this prediction, we found that the forward (*F_async_*) class was overrepresented among functional connections from layers 6, 5, 4*cβ*, 4cα, and 4A/B to layer 2/3 (Fig. 7b) (chi-squared test; 6: p<10^-5^; 5: p<10^-5^; 4cβ: p<10^-5^; 4cα: p<10^-3^; 4A/B: p<10^-5^). In contrast, functional connections within layer 2/3 exhibited a very different pattern. Within the same layer, the classes of 2/3-2/3 CCGs were more evenly represented, in stark contrast to the pattern of within-layer CCGs observed in the input layers (Fig. 6b). Within layer 2/3, the *S_sync_* and *B_sync_* CCGs were overrepresented among functional connections (Fig. 7c) (chi-squared test: p<10^-5^), and there was an equal representation of *S_sync_* and *B_sync_* CCGs among 2/3-2/3 pairings.

**Figure 7.**
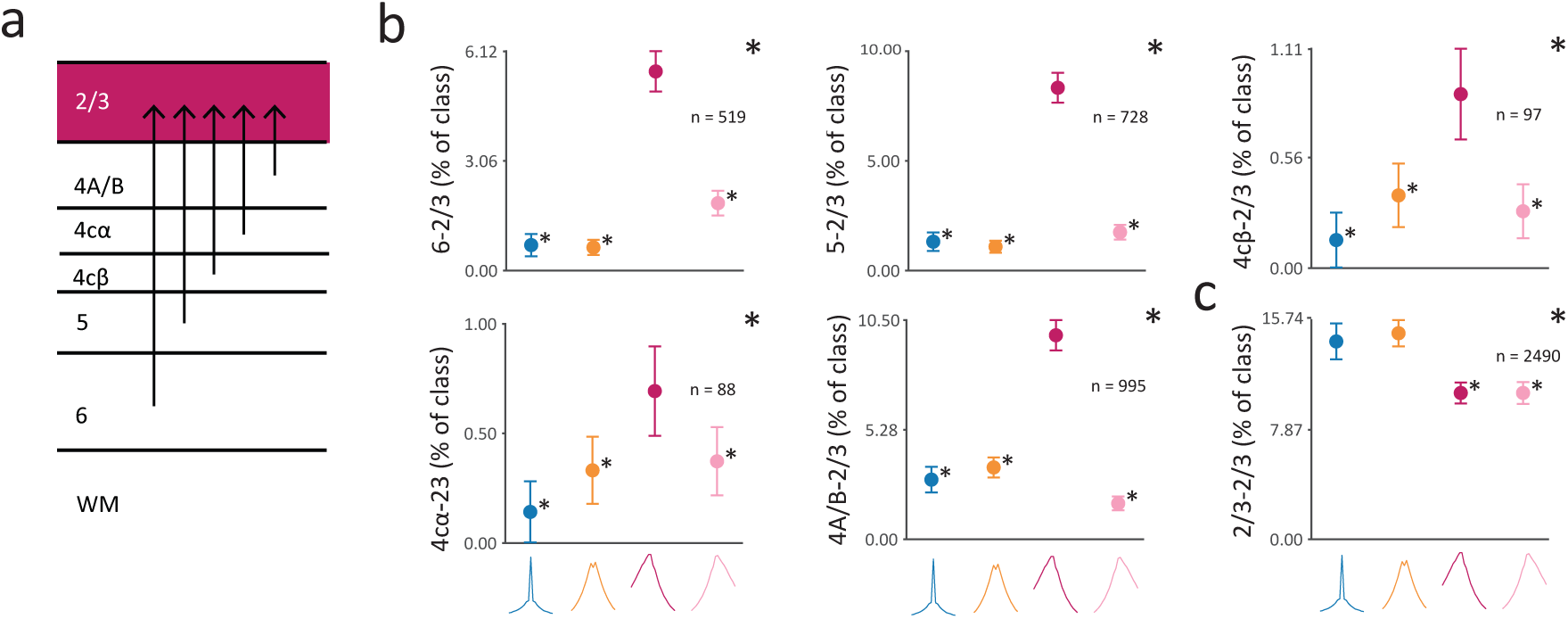
Distribution of different putative classes of functional connections with layer 2/3. **a**, Diagram of input to layer 2/3 from V1 laminar compartments. **b,** Percentage of CCGs in each class composed of one neuron in layer 2/3 and the other neuron in layer 6, 5, 4c*β*, 4cα, or 4A/B computed with the layer 2/3 neuron as the second neuron in the crosscorrelation function. **c**, Percentage of CCGs in each class composed of two neurons in layer 2/3. For b-c, conventions are the same as in Fig. 6.

## Discussion

Using high-density recordings from single neocortical columns of macaque V1, we identified 1000s of functionally connected neuronal pairs using established crosscorrelation approaches. The results demonstrate clear and systematic variations in the synchrony and strength of functional connections within single V1 columns. Notably, we observed that in spite of residing within the same column, the functional connectivity between pairs of V1 neurons depended heavily on their vertical distance within the column; both the peak lag and peak efficacy of CCGs between neuronal pairs changed dramatically within only a few hundred micrometers of vertical distance within the column. In addition, we found that the synchrony and strength of CCGs also depended on laminar location and the similarity of orientation tuning between neuronal pairs. We leveraged the statistical power provided by the large numbers of connected pairs to categorize functional connections between neurons based on their crosscorrelation functions. These analyses identified distinct classes of functional connections within the full population. Those distinct classes exhibited different distributions across defined laminar compartments, and those differences were consistent with known and/or expected properties of V1 cortical circuitry. The results demonstrate a novel utility of high-density neurophysiological recordings in assessing circuit-level interactions within local neuronal networks. Below, we discuss both the implications and the limitations of this approach.

### Effect of cortical distance on functional connectivity

A wealth of previous evidence has established a clear effect of cortical distance on functional connectivity, yet a majority of past studies have focused on the effect of horizontal distance across which large changes in shared input between neurons are expected. Evidence that spiking correlations and synchrony decline with horizontal cortical distance within V1 has been shown in cats (Das & Gilbert, 1999; Gray, Konig, Engel, & Singer, 1989; Hata, Tsumoto, Sato, & Tamura, 1991) (but see Samonds, Zhou, Bernard, & Bonds, 2006; Schwarz & Bolz, 1991), monkeys (Chu et al., 2014; Kruger & Aiple, 1988; Maldonado et al., 2000; Smith & Kohn, 2008), and in mice (Denman & Contreras, 2014). Very few studies have examined crosscorrelations among pairs of neurons within a single column, where the feedforward input is largely shared (e.g. DeAngelis et al., 1999). Longer timescale, spike count (‘noise’), correlations, which have been widely assessed in studies of primate visual cortex (Averbeck, Latham, & Pouget, 2006) have been shown to be layer dependent within macaque V1 where weaker correlations occur in layer 4 (Hansen et al., 2012). However, no evidence that such correlations depend on distance independent of layer was observed. In contrast, measurement of crosscorrelations in earlier studies of V1 columns in cat indeed suggest that functional connectivity is restricted to local regions across cortical depths (Toyama, Kimura, & Tanaka, 1981b). Within rat auditory cortex, functional connectivity diminishes dramatically within ∼300 µm of vertical columnar distance (Atencio & Schreiner, 2013), similar to what is observed in rat somatosensory cortex (Khateb, Schiller, & Schiller, 2021). A dependence of connectivity on vertical distance is further supported by evidence from multiple whole-cell recordings in mouse visual cortex which demonstrates that connection probability decreases sharply within a distance of 250 µm (Jiang et al., 2015). Our observation that the peak efficacy of CCGs was greatly diminished within <200 µm within macaque V1 is thus consistent with estimates from other sensory cortices and species.

### Distinct classes of functional connections

Upon clustering the full population of significant CCGs, we identified four putative classes of functional connections. Notably, these classes of CCGs depicted the set of pairwise connectivities that one might logically expect, namely two directional classes (forward and reverse) and two synchronous ones (sharply and broadly synchronous). More importantly, the clustering based evidence of distinct classes of CCGs was corroborated by the observation of highly differential distributions of those putative classes across cortical layers. For example, asynchronous CCG classes were more often observed among neuronal pairs within different layers, whereas synchronous pairs more often resided within the same layer. This corroboration of distinct classes extended to functional properties of V1 neurons as well in that simple cells were more often paired in a forward manner with complex cells, whereas the reverse was true for complex cells. Nevertheless, the existence of exactly four distinct classes among V1 pairs is by no means certain. Indeed, the choice of three classes was almost as valid as that of four in the clustering procedure (Fig. 4). Yet, given the clear evidence of two asynchronous classes, and the differential distributions of broadly and sharply synchronous pairs across cortical layers, the choice of only three classes of CCGs seems less parsimonious than four. Although there appeared to be less evidence for the existence of five or more classes of functional connections, that possibility cannot be ruled out either. For example, additional distinct classes of CCGs might be present, but significantly less frequent or weaker than the other four. Indeed, given their low incidence, our selection criteria already excluded CCGs with significant inhibitory peaks. As in previous studies, the frequency of excitatory CCGs in our dataset was considerably higher than that of inhibitory CCGs (Aertsen & Gerstein, 1985; Hembrook-Short et al., 2019) (Table S1).

Consequently, these additional classes were eliminated by the statistical threshold employed to identify significant CCGs. It is likely that additional distinct classes, excitatory or inhibitory, were also eliminated and/or simply fell within a mixture of the more dominant four classes that exceeded the statistical criterion. Future work will therefore be needed to more extensively characterize the distribution of distinct classes of spiking crosscorrelations among neuronal pairs in cortical columns.

### Classes of functional connections and V1 microcircuitry

We found that the four putative classes of functional connections were differentially distributed across the cortical column and across functional pairs of neurons. Most notably, the different CCG classes were observed in different proportions across V1 layers. In spite of those differences however, it need not follow that the relative proportion of any specific putative class (e.g. sharply synchronous) fits with the known (or predicted) connectivity between different V1 neurons. For example, our observation that asynchronous CCGs (*F_async_* and *R_async_*) were considerably more frequent among neuronal pairs situated in different laminae, and that synchronous CCGs (*S_sync_* and *B_sync_*) were found among neurons in the same laminae, would not be expected if the differences in synchrony resulted primarily from measurement noise. Likewise, the observed disproportionality of *F_async_* and *R_async_* CCGs among functional connections between simple and complex cells would not be expected if the two asynchronous classes were indistinguishable in our measurements. Instead, not only were the two classes disproportionate among simple and complex cells, but the overall direction of disproportionality was consistent with the known connectivity between the two functional classes of cells (Alonso & Martinez, 1998; Martinez & Alonso, 2001; Yu & Ferster, 2013).

Overall, we found that the pattern of differential distributions of the putative classes of CCGs across the column and across functional pairs of neurons was largely consistent with known properties of V1 microcircuitry. Nevertheless, it is important to emphasize that spike crosscorrelations are in no way a substitute for more direct measures of synaptic connectivity, e.g. multi-patch recordings (Jiang et al., 2015). Instead, the relative instances of different types of crosscorrelations observed among large populations of neuronal pairs, as shown here, provide a means of constraining models of cortical microcircuits. This approach should prove particularly valuable in less experimentally tractable model systems such as nonhuman primates, or perhaps even in the human brain, where direct measurements are not yet possible. In such cases, the ability of high-channel count, high-density, probes to dramatically increase the number of identifiable functional connections within a local network of neurons is among their greater benefits. Our results thus far suggest that this approach works well and could be extended to examine higher-order connectivities among larger sets of neurons, and to identify neuronal ensembles with distinct functional properties (Fujisawa, Amarasingham, Harrison, & Buzsaki, 2008; Miller, Ayzenshtat, Carrillo-Reid, & Yuste, 2014; See, Atencio, Sohal, & Schreiner, 2018). In addition, future studies should be able to compare local connectivities across different putative cell types estimated from their spike waveforms (Johnston, DeSouza, & Everling, 2009; E. K. Lee et al., 2021; Mitchell, Sundberg, & Reynolds, 2007; Wilson, O’Scalaidhe, & Goldman-Rakic, 1994) and/or spiking patterns (Onorato et al., 2020). Combined with measurements of functional connectivity, such an approach could be used to constrain models of microcircuit architecture from neurophysiological data obtained from any number of uniquely evolved primate brain structures.

## Acknowledgments

We thank Jonathan C. Horton for extensive help with the recordings and histology. We thank Tim Harris and Karel Svoboda for providing the Neuropixel probes, Shellie Hyde and Sam Baker for technical assistance. This work was supported by NIH Grant EY014924 and EY029759.

## Author Contributions

S.Z., X.C. and T.M. designed the research and performed the experiments. E.T., S.Z., and R.X. analyzed the data. E.T., S.Z. and T.M. wrote the paper.

## Data Availability

All of the raw data collected in study will be made publicly accessible and available for further analyses at request.

## Methods

### Experimental Model and Subject Details

Anesthetized recordings were conducted in 2 adult male macaques (M1, 13 kg; M2 8 kg). All experimental procedures were in accordance with National Institutes of Health Guide for the Care and Use of Laboratory Animals, the Society for Neuroscience Guidelines and Policies, and Stanford University Animal Care and Use Committee.

### Electrophysiological Recordings

Prior to each recording session, treatment with dexamethasone phosphate (2 mg per 24 h) was instituted 24 h to reduce cerebral edema. After administration of ketamine HCl (10 mg per kilogram body weight, intramuscularly), monkeys were ventilated with 0.5% isoflurane in a 1:1 mixture of N2O and O2 to maintain general anesthesia. Electrocardiogram, respiratory rate, body temperature, blood oxygenation, end-tidal CO2, urine output and inspired/expired concentrations of anesthetic gases were monitored continuously. Normal saline was given intravenously at a variable rate to maintain adequate urine output. After a cycloplegic agent was administered, the eyes were focused with contact lenses on a CRT monitor. Vecuronium bromide (60 µg/kg/h) was infused to prevent eye movements.

With the anesthetized monkey in the stereotaxic frame, an occipital craniotomy was performed over the opercular surface of V1. The dura was reflected to expose a small (∼3 mm^2^) patch of cortex. Next, a region relatively devoid of large surface vessels was selected for implantation, and the Neuropixels probe was inserted with the aid of a surgical microscope. Given the width of the probe (70 um x 20 um), insertion of it into the cortex sometimes required multiple attempts if it flexed upon contacting the pia. The junction of the probe tip and the pia could be visualized via the (Zeiss) surgical scope and the relaxation of pia dimpling was used to indicate penetration, after which the probe was lowered at least 3-4 mm. Prior to probe insertion, it was dipped in a solution of the DiI derivative FM1-43FX (Molecular Probes, Inc) for subsequent histological visualization of the electrode track.

Given the length of the probe (1 cm), and the complete distribution of electrode contacts throughout its length, recordings could be made either in the opercular surface cortex (M1) or within the underlying calcarine sulcus (M2), by selecting a subset of contiguous set of active contacts (n = 384) from the total number (n=986). Receptive fields (RFs) from online multi-unit activity were localized on the display using at least one eye. RF eccentricities were ∼ 4-6° (M1) and ∼ 6-10° (M2). Recordings were made at 1 to 3 sites in one hemisphere of each monkey. At the end of the experiment, monkeys were euthanized with pentobarbital (150 mg kg^−1^) and perfused with normal saline followed by 1 liter of 1% (wt/vol) paraformaldehyde in 0.1 M phosphate buffer, pH 7.4.

### Visual Stimulation

Visual stimuli were presented on a LCD monitor NEC-4010 (Dimensions= 88.5 (H)* 49.7 (V) cm, pixels=1360 * 768, frame rate= 60 Hz) positioned 114 cm from the monkey. Stimuli consisted of circular drifting Gabor gratings (2 deg./sec., 100% Michelson contrast) positioned within the joint RFs of recorded neurons monocularly. Gratings drifted in 36 different directions between 0 to 360° in 10° steps in a pseudorandom order. Four spatial frequencies (0.5, 1, 2, 4 cycle/deg.) were tested and optimal SFs were determined offline for data analysis. The stimulus in each condition was presented for 1 second and repeated 5 or 10 times. A blank screen with equal luminance to the Gabor patch was presented for 0.25s during the stimulus interval.

### Layer Assignment

The laminar location of our recording sites was estimated based on a combination of functional analysis and histology results. For each recording, we first performed the current source density (CSD) analysis on the stimulus-triggered average of local field potentials (LFP). LFP were low-pass filtered at 200 Hz and recorded at 2500 Hz. LFP signals recorded from each 4 neighboring channels were averaged and realigned to the onset of visual stimulus. CSD was estimated as the second-order derivatives of signals along the probe axis using the common five-point formula (Nicholson & Freeman, 1975). The result was then smoothed across space (σ = 120 µm) to reduce the artifact caused by varied electrode impedance. We located the lower boundary of the major sink (the reversal point of sink and source) as the border between layer 4C and layer 5/6. Based on this anchor point, we assign other laminar compartment borders using the histological estimates.

### Single neuron properties

To characterize the visual properties of each neuron, the stimulus evoked activity was assessed using mean firing rate (spikes/sec) over the entire stimulus presentation period, offset by a response latency of 30 ms. Only responses to the preferred spatial frequency were used. Modulation ratio was defined as F1/F0, where F1 and F0 are the amplitude of the first harmonic at the temporal frequency of drifting grating and constant component of the Fourier spectrum to the neuron’s response to preferred orientation. Simple cells were defined as cells with modulation ratio larger than 1, and complex cells have modulation ratios smaller than 1 (De Valois et al., 1982; Skottun et al., 1991).

### Signal Correlations

To measure the similarity of orientation tuning between neuronal pairs, we computed an orientation signal correlation (r_ori_). The orientation signal correlation was defined as the Pearson’s correlation coefficient between the mean responses of two neurons to each of the 36 stimulus orientations (Smith & Kohn, 2008). For each neuron and orientation, a single mean response was computed by averaging spiking activity over the entire duration of stimulus presentation (1 second) across all trials with a particular orientation.

### Cross-correlograms (CCGs)

To measure correlated firing, we computed the crosscorrelation between spike trains of all pairs of simultaneously recorded neurons (Jia et al., 2013; Siegle et al., 2021; Smith & Kohn, 2008; Zandvakili & Kohn, 2015). We focused on the spiking activity within the 0 second window of each visual stimulus presentation, which ensured that the analysis was not affected by the transient response to stimulus onset. To mitigate firing rate effects, we normalized the crosscorrelation for each pair of neurons by the geometric mean of their firing rates. Thus, the cross-correlogram *CCG* for a pair of neurons (*j*, *k*)was defined as follows:

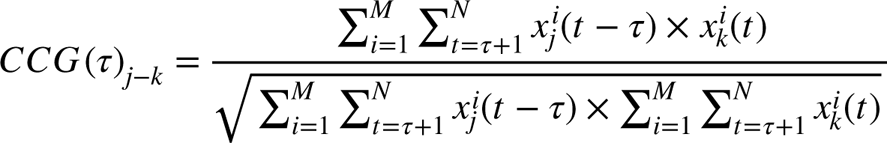

where is *M* the number of trials, *N* is the number of time bins within a trial, *τ* is the time lag, and *x^i^_j_*(*t*) is one if neuron *j* fired in time bin of trial and zero otherwise. We denote the CCG computed with neuron j as the first neuron in the correlation function as j-k and the CCG computed with neuron k as the first neuron as k-j.

To correct for correlation due to stimulus-locking or slow fluctuations in population response (e.g., gamma-band activity), we computed a jitter-corrected cross-correlogram by subtracting a jittered cross-correlogram from the original cross-correlogram:

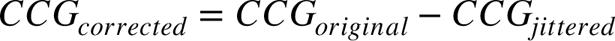

The jittered cross-correlogram (*CCG_jittered_*) reflects the expected value of cross-correlograms computed from all possible jitters of each spike train within a given jitter window (Harrison & Geman, 2009; Smith & Kohn, 2008). The jittered spike train preserves both the PSTH of the original spike train across trials and the spike count in the jitter window within each trial. As a result, jitter correction removes the correlation between PSTHs (stimulus-locking) and correlation on timescales longer than the jitter window (slow population correlations). Here, a 25-ms jitter window was chosen based on previous studies (Jia et al., 2013; Siegle et al., 2021; Zandvakili & Kohn, 2015).

We classified a CCG as significant if the peak of the jitter-corrected CCG occurred within ms of zero and was more than seven standard deviations above the mean of the noise distribution. The noise distribution for a CCG was defined as the flanks of the jittered-corrected CCG ({*CCG* (*τ*) | 100 ≥ |*τ*| ≥ 50}). This significance criterion was chosen based on that of Siegle et al. 2021. All analyses presented here involve only significant, jitter-corrected cross-correlograms. Note that the criterion identifies only positive peaks in the CCG and excludes significant inhibitory correlations. However, consistent with earlier studies (Aertsen & Gerstein, 1985; Hembrook-Short et al., 2019), we found that the frequency of CCGs with significant troughs was approximately 40x lower than those with significant peaks (Table S1).

### Classification of Cross-correlogram

To identify distinct classes of crosscorrelation functions, we clustered significant crosscorrelations. We only analyzed crosscorrelation functions between *τ* = −10 and *τ* = 10 such that our input CCGs had 21 features, corresponding to the 21 crosscorrelation values between *τ* = -10 and *τ* = 10. For clustering, we included two crosscorrelation functions for each pair of neurons ( *j*, *k*), one computed using the above CCG function with neuron j as the first neuron *j* – *k* and the other with neuron k as the first neuron (*k* − *j*). This corresponds to reflecting the CCG function across the vertical line *τ* = 0. We z-scored each input CCG prior to clustering to encourage clustering based on the shape of the correlation function rather than its magnitude.

To simplify the clustering problem, we used t-distributed stochastic neighbor embedding (t-SNE) to reduce our input data with 21 features to 3 features (tsne, MATLAB R2019a). t-SNE was used instead of principal component analysis (PCA) because it is more robust to outliers since it captures neighbor relationships in the input space. We clustered the dimensionality-reduced data using k-means with *k* to (50 replicates, 100 max iterations, kmeans MATLAB R2019a). To determine the optimal number of clusters, we used two complementary approaches, the elbow method and silhouette method. The elbow method selects *k* based on the magnitude of the change in the variance explained by clustering as *k* increases. For a set of points *S* = _{_*s*_1_, *s*_2_, …, *s_n_*_}_ divided into *k* clusters *S*_1_, *S*_2_, …, *S_k_*, the percent of variance explained by clustering (*η_k_*) is:

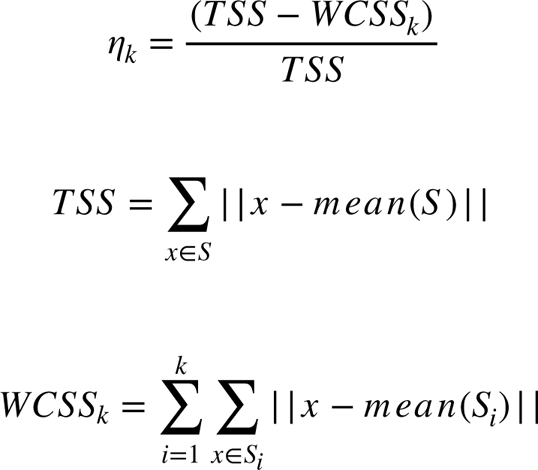

where *TSS* denotes the total sums of squares and *WCSS_k_* denotes the sum of within cluster sums of squares over all clusters. The optimal number of clusters occurs at the point where the percent of explained variance plateaus (or ‘elbows’) as the number of clusters increases. The silhouette criterion captures how similar a point is to its own cluster versus how different it is from the nearest cluster that it is not a member of. We computed the silhouette criterion using MATLAB’s evalclusters function with default parameters (MATLAB R2019a).

### Statistical Analyses

The effects of vertical pair distance and orientation signal correlation on CCG peak lag and peak efficacy were fit using linear and exponential functions. In linear regressions predicting CCG peak lag, all significant CCGs were included, and mean squared error was used as the cost function for regressions. In linear and exponential regressions predicting CCG peak efficacy, only significant CCGs with non-outlier peaks (1.5*IQR criterion) were included, and mean absolute error was used as the cost function for regressions to encourage fit of the plotted median peak efficacies.

The relationships between classes of functional connections and signal correlation/pair distance were evaluated using Wilcoxon rank-sum tests, and the relationship between functional class and layer/cell type pairings was assessed using chi-squared tests or one proportion z-tests. Finally, the dependence of functional class on whether a CCG was composed of two neurons within the same or different cortical layer(s) with comparable vertical distance was assessed using logistic regression.

### Distance matching

Distance matching was used to compare the effects of orientation signal correlation on CCG peak lag and peak efficacy among neuronal pairs with comparable cortical distances. To match pairs with comparable distances (Fig. 3C), we sorted significant CCGs by cortical distance, then paired the CCGs with the smallest and second smallest distances and paired the CCGs with the third and fourth smallest distances and so forth. Thus, every significant CCG was paired with exactly one other significant CCG, resulting in 5,122 pairs. To verify that this procedure effectively matched pairs of CCGs with comparable cortical distance, we examined the difference in cortical distance for distance-matched pairs. More than 99% (5,067/5,122) of the distance-matched pairs had a difference in cortical distance of less than 2 µm. Finally, we examined the correlation between the difference in CCG peak lag or peak efficacy and difference in signal correlation for matched pairs to determine whether signal correlation predicts peak lag or peak efficacy when controlling for distance.

---

**Figure S1.**
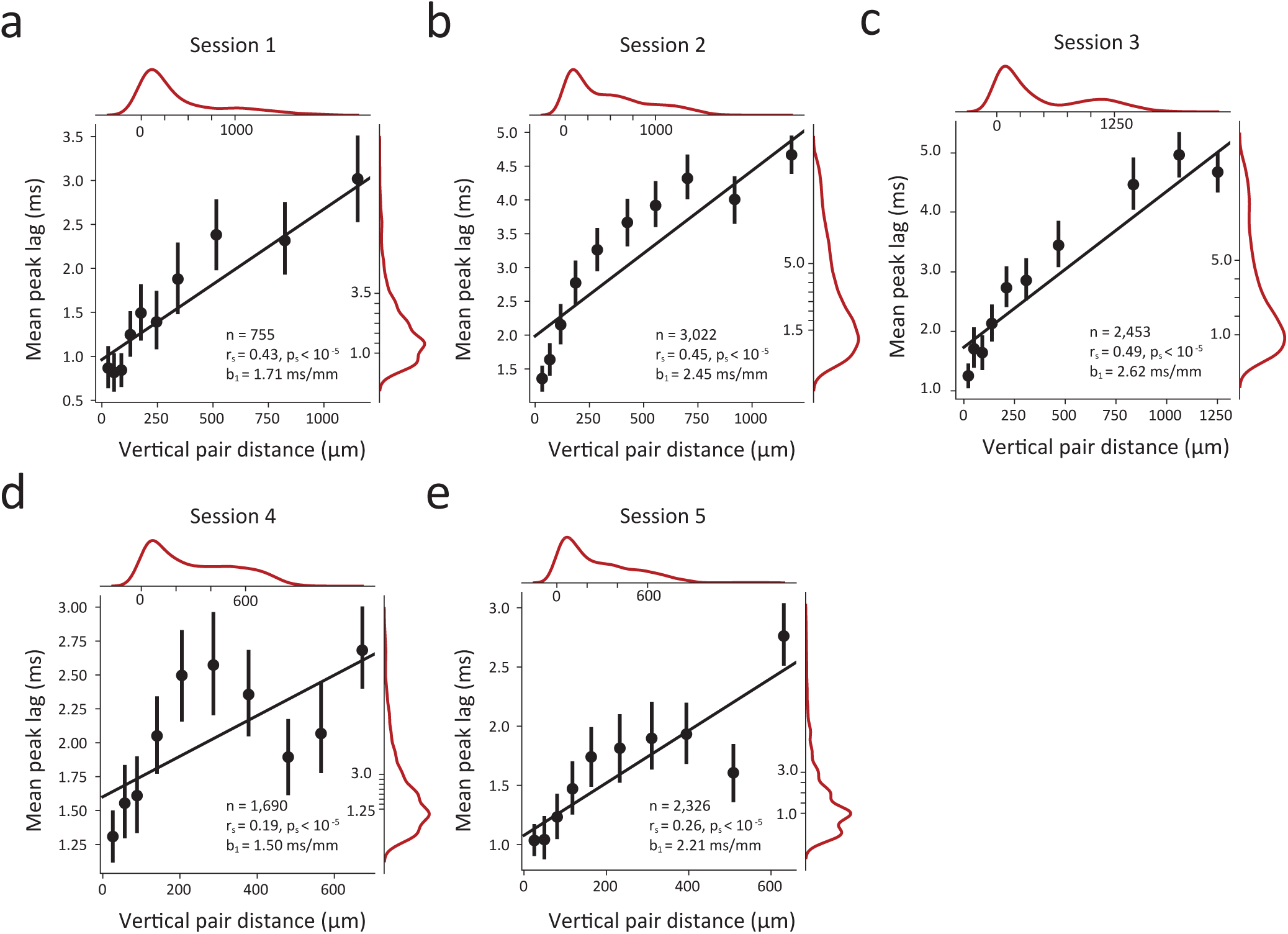
Dependence of synchrony on vertical distance within single cortical columns for all individual sessions. Linear dependence of synchrony on vertical pair distance for each of the 5 sessions (a-e). Plots follow the conventions used in Figure 2.

**Figure S2.**
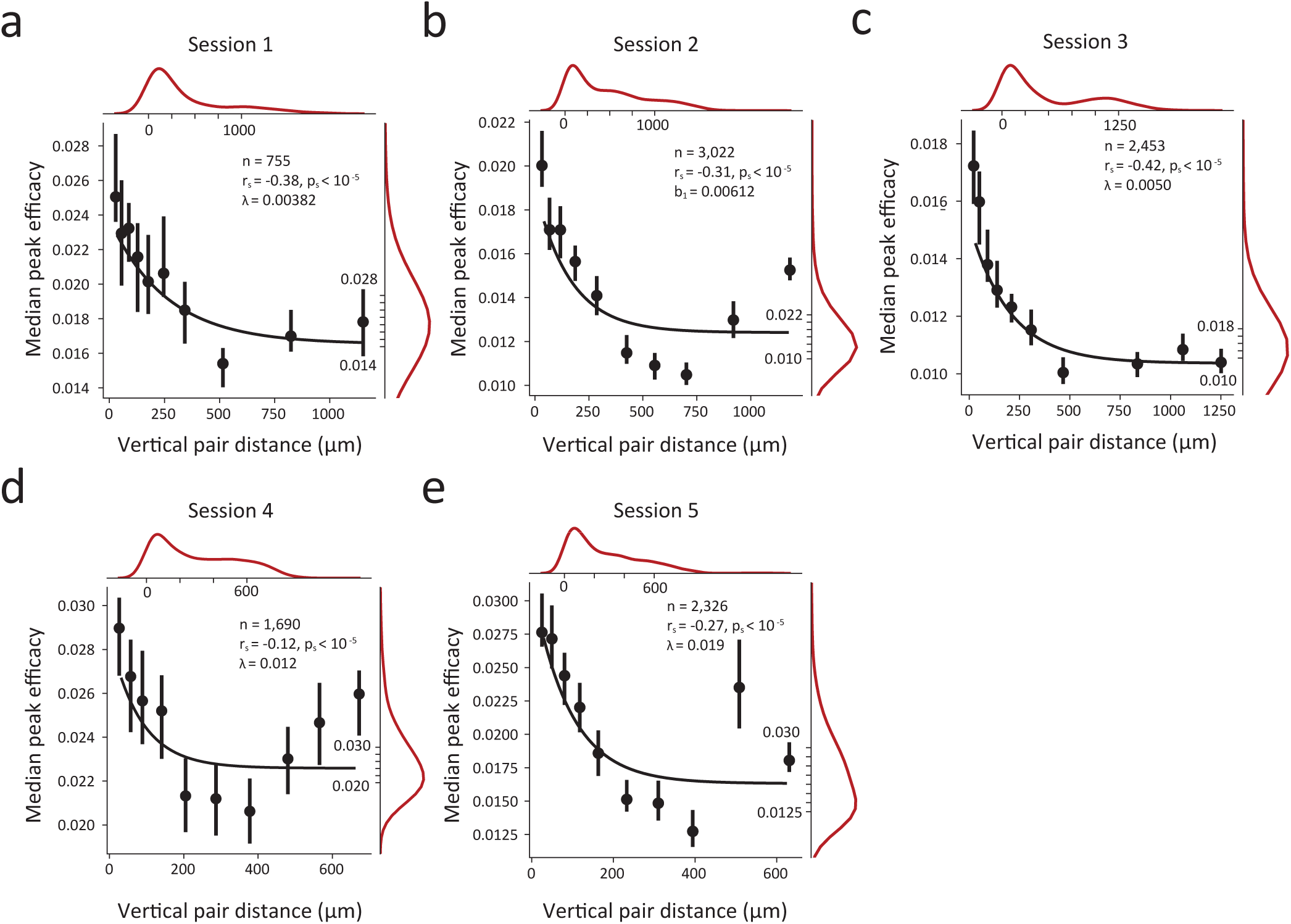
Dependence of the strength of functional connectivity on vertical distance within single cortical columns for all individual sessions. Peak efficacy of CCGs decays with greater pair distance in each of the 5 sessions (a-e). Plots follow the conventions used in Figure 2.

**Figure S3.**
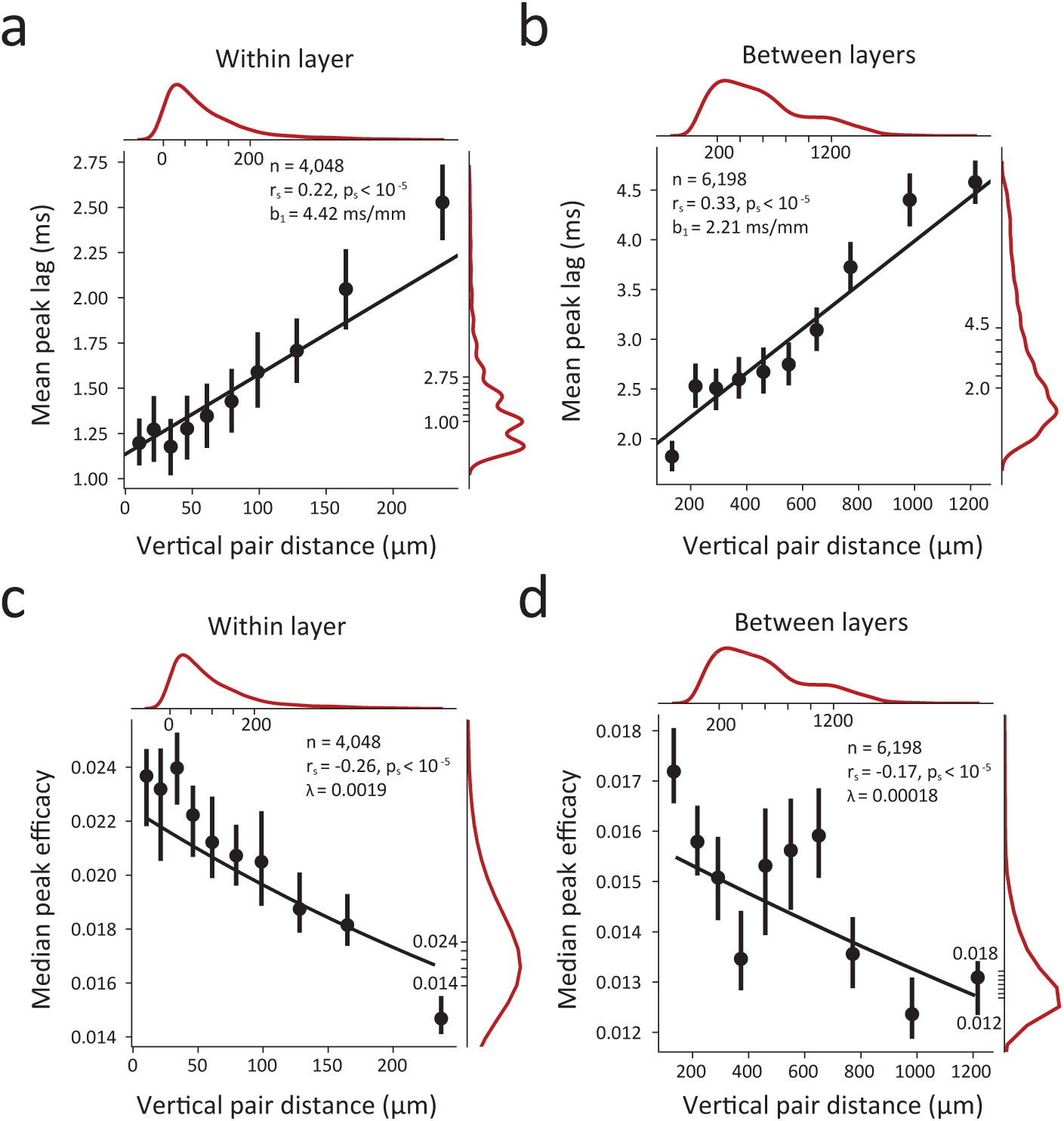
Dependence of the synchrony and strength of functional connectivity on vertical distance within single cortical columns, separated by laminar pairings. (a-b) Linear dependence of CCG peak lag on the distance between neurons for pairs of neurons composed of two neurons in the same layer of V1 (a) or two neurons in different layers of V1 (b). (c-d) Peak efficacy of CCGs decays with greater pair distance in pairs of neurons composed of two neurons in the same layer (c) and different layers (d). Plots follow the conventions used in Figure 2.

**Figure S4.**
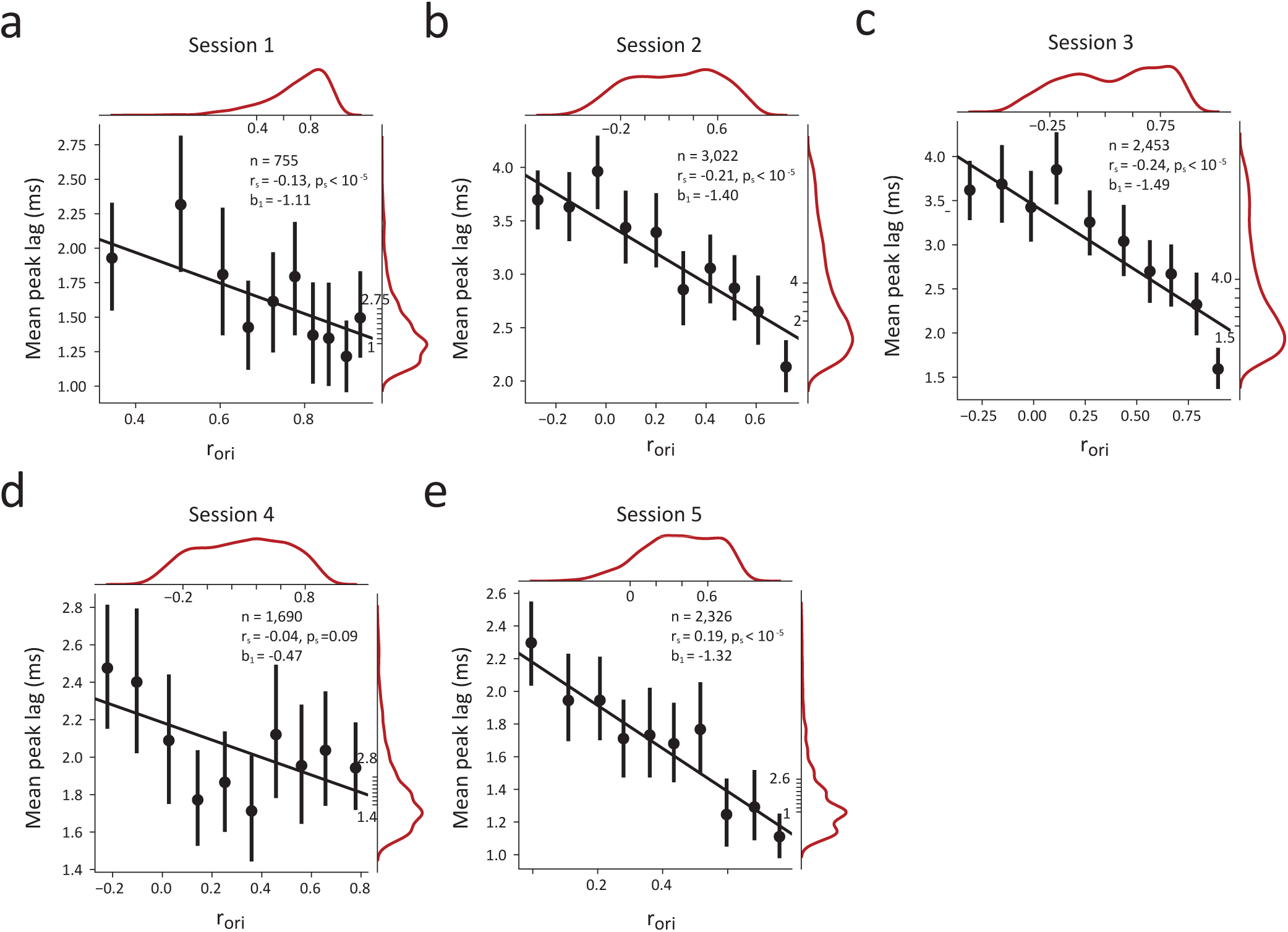
Dependence of synchrony on tuning similarity within single cortical columns for all individual sessions. Linear dependence of synchrony on tuning similarity in each of the 5 sessions (a-e). Plots follow the conventions used in Figure 3.

**Figure S5.**
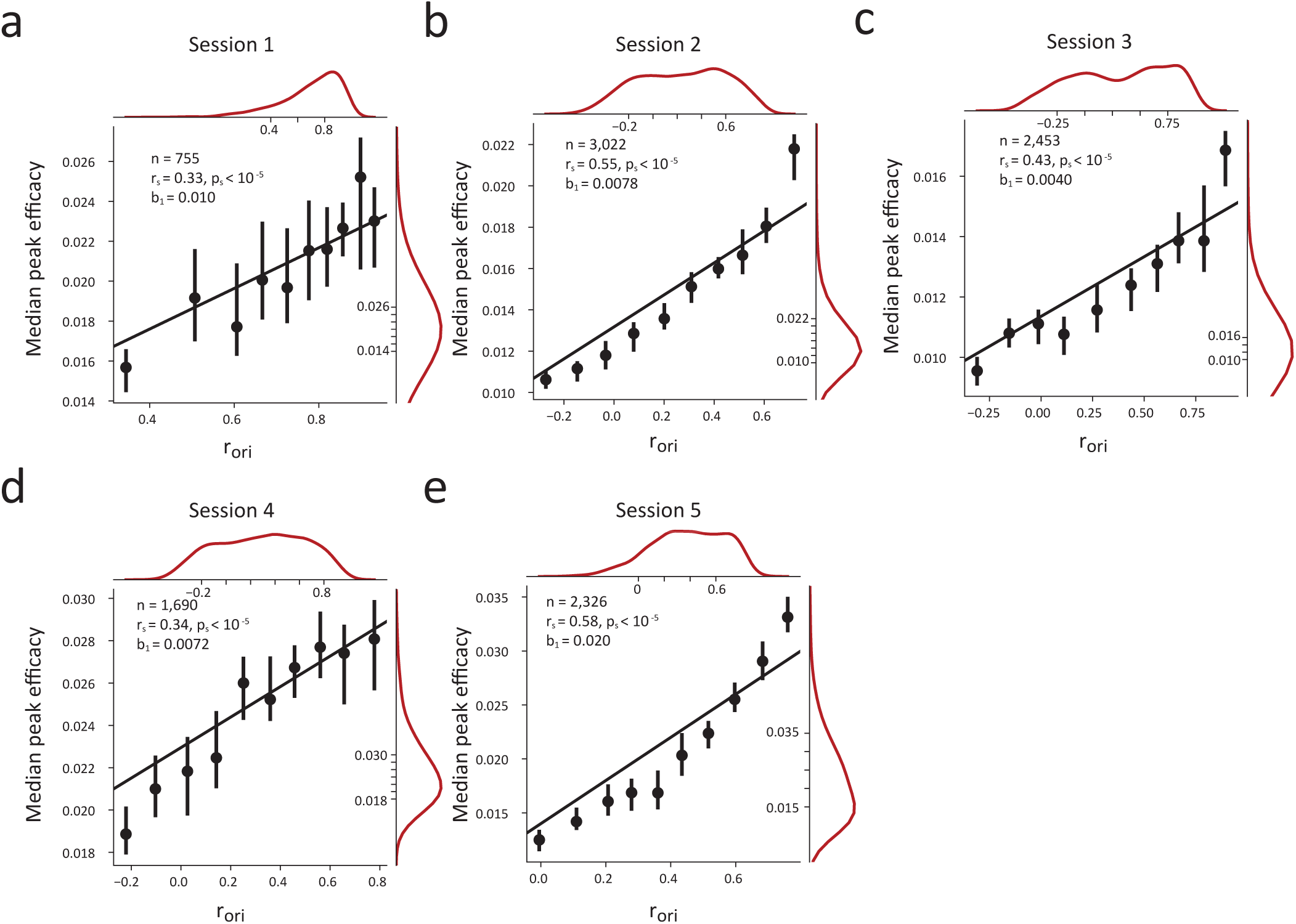
Dependence of the strength of functional connectivity on tuning similarity within single cortical columns for all individual sessions. Peak efficacy of CCGs is positively correlated with tuning similarity in each of the 5 sessions (a-e). Plots follow the conventions used in Figure 3.

**Figure S6.**
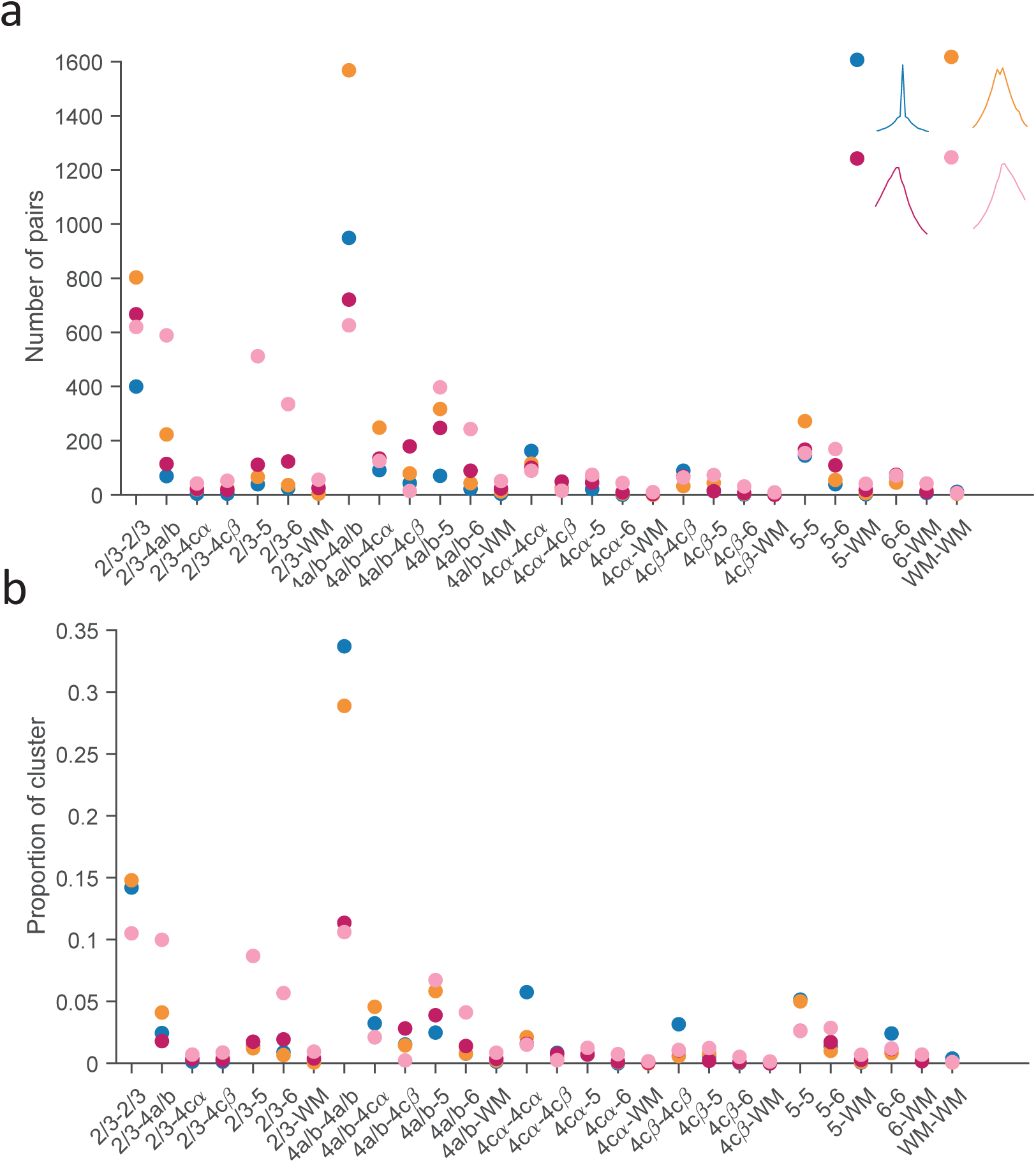
Distribution of putative CCG classes across layer pairings. Plotted is the number (a) and proportion (b) of significant CCGs in each class composed of two neurons in the indicated layer pairing. For example, CCGs in the 2/3-4a/b pairing are composed of one neuron in layer 2/3 and one in 4a/b and are computed with the 2/3 neuron first in the crosscorrelation function.

**Table S1.** Number of CCGs with peaks or troughs significantly above or below noise. The number of recorded pairs with a peak or trough at least 1-7 standard deviations above or below noise is shown. Only CCGs with peaks or troughs within 10 ms of zero time lag were considered. 136 CCGs had troughs that were more than 7 SD below noise whereas 10,246 CCGs had peaks that were more than 7 SD above noise.

| **Stdev.**<br>**Above/Below**<br>**Noise** | **Number of**<br>**Pairs with**<br>**Peaks Above**<br>**Noise** | **Number of**<br>**Pairs with**<br>**Troughs Below**<br>**Noise** |
| --- | --- | --- |
| **1** | 33,502 | 15,582 |
| **2** | 33,502 | 11,347 |
| **3** | 31,660 | 4,537 |
| **4** | 25,180 | 1,555 |
| **5** | 18,739 | 603 |
| **6** | 13,757 | 265 |
| **7** | 10,246 | 136 |

**Table S2.** Dependence of putative classes on laminar pairing and vertical distance for pairs of neurons separated by 86-310 μm. Coefficients, standard errors, and p-values from logistic regressions predicting class membership using the distance between pairs of neurons and whether pairs were located in the same or different cortical layer(s). Only pairs of neurons with pair distances greater than the 5% of pairs located in different cortical layers (>86 μm) and less than 5% of pairs located in the same cortical layer (<310 μm) were included. Significant predictors (p<10^-5^) are highlighted.

| Dependent Variable<br>(in/out of cluster) | Predictor | Coefficient | Standard Error | P-Value |
| --- | --- | --- | --- | --- |
| $S_{sync}$ | Distance | -0.0041/$\mu\text{m}$ | 0.0007 | **$2.78 \times 10^{-8}$** |
|  | Layer | 0.060/layer | 0.090 | 0.50 |
| $B_{sync}$ | Distance | -0.0017/$\mu\text{m}$ | 0.0005 | $3.53 \times 10^{-4}$ |
|  | Layer | 0.280/layer | 0.060 | **$3.18 \times 10^{-6}$** |
| $F_{async}$ | Distance | 0.0016/$\mu\text{m}$ | 0.0005 | $9.35 \times 10^{-4}$ |
| | Layer | -0.172/layer | 0.0615 | $5.24 \times 10^{-3}$ |
| $R_{async}$ | Distance | 0.0021/$\mu\text{m}$ | 0.0005 | $2.53 \times 10^{-5}$ |
|  | Layer | -0.160/layer | 0.0631 | 0.011 |

